# Natural malaria infection elicits rare but potent neutralizing antibodies to the blood-stage antigen RH5

**DOI:** 10.1101/2023.10.04.560669

**Authors:** Lawrence T. Wang, Andrew J.R. Cooper, Brendan Farrell, Kazutoyo Miura, Ababacar Diouf, Nicole Müller-Sienerth, Cécile Crosnier, Lauren Purser, Maciej Maciuszek, Jordan R. Barrett, Kirsty McHugh, Courtney Tucker, Shanping Li, Safiatou Doumbo, Didier Doumtabe, Chul-Woo Pyo, Carolyn M. Nielsen, Sarah E. Silk, Kassoum Kayentao, Aissata Ongoiba, Ming Zhao, Doan C. Nguyen, F. Eun-Hyung Lee, Angela M. Minassian, Daniel E. Geraghty, Boubacar Traore, Robert A. Seder, Peter D. Crompton, Gavin J. Wright, Carole A. Long, Simon J. Draper, Matthew K. Higgins, Joshua Tan

## Abstract

*Plasmodium falciparum* RH5 is the most advanced blood-stage malaria vaccine candidate and is under evaluation for efficacy in endemic regions, emphasizing the need to study the underlying antibody response to RH5 during natural infection. Here, we found that RH5-reactive B cells were rare in malaria-exposed individuals despite repeated infections over multiple years. RH5-specific monoclonal antibodies isolated from these individuals were extensively mutated but mostly targeted non-neutralizing epitopes, in contrast to antibodies from RH5-vaccinated, malaria-naive individuals. However, infection-derived MAD8-151 and MAD8-502 were among the most potent neutralizers out of 186 antibodies isolated from both cohorts and target the same epitopes as the most effective vaccine-induced antibodies. Binding to basigin receptor-proximal epitopes was the primary factor governing the potency of RH5-specific antibodies from both natural infection and vaccination, followed by the strength of binding. These results indicate a clear strategy for the development of next-generation RH5 vaccines for use in malaria-endemic regions.

## INTRODUCTION

Developing highly effective tools against *Plasmodium falciparum*, the parasite that causes the most severe form of malaria, is a long-sought global health priority (WHO, 2021). Encouraging progress has been made recently in the development of vaccines and monoclonal antibodies (mAbs) that target *P. falciparum* sporozoites, the parasite stage transmitted by *Anopheles* mosquitoes to initiate infection in the human host (Datoo et al., 2022; Kayentao et al., 2022; Mwakingwe-Omari et al., 2021; Wu et al., 2022). However, the protection provided by these interventions does not extend to the subsequent disease-causing blood stage of infection due to antigenic differences between the two parasite stages. Given the potential for only a single breakthrough sporozoite to cause malaria disease, there is a strong impetus to develop tools and strategies that target blood-stage parasites as a second line of defense. However, decades of effort to develop blood-stage vaccines have been hampered by the parasite’s ability to evade antibodies by mutating its surface antigens and invading erythrocytes via multiple redundant pathways (Takala et al., 2009; Wright and Rayner, 2014).

Blood-stage vaccine development was bolstered by the identification of a well-conserved tripartite complex used by *P. falciparum* merozoites to invade host erythrocytes. This complex was initially found to be composed of reticulocyte-binding protein homologue 5 (RH5), cysteine-rich protective antigen (CyRPA), and RH5-interacting protein (Ripr) (Ragotte et al., 2020; Wong et al., 2019). A recent study identified two other binding partners in this complex: *Plasmodium* thrombospondin-related apical merozoite protein (PTRAMP) and cysteine-rich small secreted protein (CSS) (Knuepfer et al., 2019; Scally et al., 2022). Of all the members of this complex, RH5 is the most well-studied as a vaccine candidate and is at the most advanced stage of clinical trials (Silk et al., 2023). This antigen plays an indispensable role in mediating merozoite invasion by binding the erythrocyte surface protein basigin (Baum et al., 2009; Crosnier et al., 2011; Wright et al., 2014). Furthermore, RH5 vaccination can elicit broadly neutralizing antibodies in animals (Bustamante et al., 2013; Douglas et al., 2011) and a single RH5-specific mAb conferred protection against blood-stage *P. falciparum* challenge in *Aotus* monkeys (Douglas et al., 2019). Nevertheless, the biological role and location of RH5 during erythrocyte invasion present unique challenges for vaccine development. RH5 is not constitutively expressed on the merozoite surface but is sequestered within intracellular organelles and is only released to the surface just prior to engagement of basigin, providing a limited time window for antibodies to bind. Therefore, antibodies targeting RH5 need to be present at sufficient concentrations and must bind quickly to prevent merozoite invasion. This challenge is illustrated by findings from a recent Phase I/II clinical trial, in which malaria-naïve individuals were vaccinated with a recombinant RH5 vaccine and subsequently exposed to controlled human malaria infection (CHMI) (Minassian et al., 2021). This trial reported that the vaccine significantly delayed blood-stage growth of parasites but did not elicit sufficient neutralizing antibodies to prevent malaria diagnosis (Minassian *et al*., 2021). Therefore, more work is needed to improve the ability of RH5 vaccines to elicit both higher antibody titers and a more focused response towards regions of the protein that interfere with merozoite function.

A key recent development in the RH5 vaccine field is the initiation of clinical trials in sub-Saharan Africa to test RH5 vaccine safety, immunogenicity, and efficacy against clinical malaria in endemic regions (Silk *et al*., 2023). However, the longevity and function of underlying anti-RH5 antibodies and the frequency of RH5-reactive B cells elicited by natural infection remain poorly understood, aside from reports that anti-RH5 polyclonal antibody titers in malaria-exposed humans are low (Douglas *et al*., 2011; Partey et al., 2018; Tran et al., 2014; Willcox et al., 2021) but correlate with protection against malaria (Chiu et al., 2014; Healer et al., 2019; Richards et al., 2013; Tran *et al*., 2014). More generally, much remains to be elucidated about the molecular details of the human antibody response to RH5. To our knowledge, only a single study on human RH5-specific mAbs has been reported, in which 17 mAbs were isolated from malaria-naïve, RH5-vaccinated individuals and characterized in detail (Alanine et al., 2019). Furthermore, no such studies of RH5-specific antibodies from naturally infected individuals have been performed.

Here, we investigated the B cell response to RH5 during natural malaria infection and compared this to the response elicited by RH5 vaccination. We found evidence of a weak response to RH5 despite repeated malaria infections over multiple years, both at the level of RH5-reactive B cells and circulating anti-RH5 antibodies. We analyzed the binding strength, specificity, and function of a panel of 186 mAbs derived from natural infection and RH5 vaccination. Neutralization potency was strongly associated with binding to three distinct regions of RH5 proximal to the receptor-binding site that contacts basigin, which we refer to as the “top” of the protein. While mAbs induced by malaria infection were on average less potent than mAbs derived from vaccination, two infection-derived mAbs (MAD8-151 and MAD8-502) targeted critical RH5 epitopes and were among the most potently neutralizing mAbs in the entire panel. Moreover, we found evidence for shared V gene usage between infection- and vaccination-derived mAbs. In particular, infection-derived MAD8-151 and vaccination-derived MAD10-255 targeted the same epitope with similar potency and shared identical heavy and light chain V(D)J gene usage and CDR1-3 lengths, providing evidence for convergent B cell selection despite differences in the type of exposure to RH5 and donor geographic origin. These findings highlight vulnerable regions of RH5 targeted by potently neutralizing antibodies and suggest that natural infection is capable of inducing neutralizing antibody responses that can potentially be boosted by vaccination, providing valuable insight for fine-tuning next-generation RH5 vaccines.

## RESULTS

### Malaria exposure elicits weak B cell responses to RH5 despite repeated infection

To investigate the antibody response to RH5 during natural infection, we studied a longitudinal cohort of 758 individuals living in Kalifabougou, Mali (Tran et al., 2019). This rural community experiences intense seasonal malaria between July and December, during which the estimated entomological inoculation rate is 2 infective bites per person per day at the peak of the season. Annual collection of samples before and after the malaria season, as well as during active and passive surveillance for clinical malaria cases, allowed us to study antibody levels to RH5 during critical time points. To analyze long-lasting antibody responses to *P. falciparum* antigens, we initially focused on plasma collected in May after the five-month dry season when malaria transmission is minimal. RH5-specific IgG levels in plasma from all 758 donors were compared with IgG levels to merozoite surface protein-1 (MSP1), an immunodominant protein expressed during the same stage of the parasite life cycle (Obeng-Adjei et al., 2020; Tran *et al*., 2014). Polyclonal IgG binding was lower for RH5 compared to MSP1 (23.1% versus 95.1% individuals with binding above background, respectively) (Figure 1A). We examined whether IgG responses to RH5 and MSP1 were associated with age in a cross-sectional survey (Figure 1B). MSP1-specific IgG levels increased with age, particularly in the first 5 years, consistent with rapid activation of B cells by an immunodominant blood-stage antigen. However, there was no association between age and RH5-specific IgG levels, suggesting that repeated exposure to merozoites over multiple malaria seasons did not trigger a long-lasting antibody response to this antigen. This could either be due to the inability of RH5 to activate B cells or the induction of only short-lived antibody-secreting B cells during infection. To distinguish between the two possibilities, we screened paired acute (collected on the day of malaria diagnosis) and convalescent (∼1 week after acute sample collection) plasma samples from donors who had documented cases of clinical malaria. Antibody levels to MSP1 increased in this one-week window (*P* < 0.01), confirming that these samples captured a suitable time period to measure the induction of anti-merozoite responses (Figure 1C). Antibody levels to RH5 also increased in this time period (P < 0.01), indicating that natural infection triggers a detectable antibody response to RH5. Indeed, when we tracked the RH5-specific IgG profile of 4 individuals over a period of 6 years, we observed frequent occasions in which a documented case of clinical malaria resulted in a sharp increase in RH5 antibody levels (Figure 1D). However, these were often followed by a rapid decline, consistent with the induction of only short-lived antibody responses against RH5 after infection.

**Figure 1:**
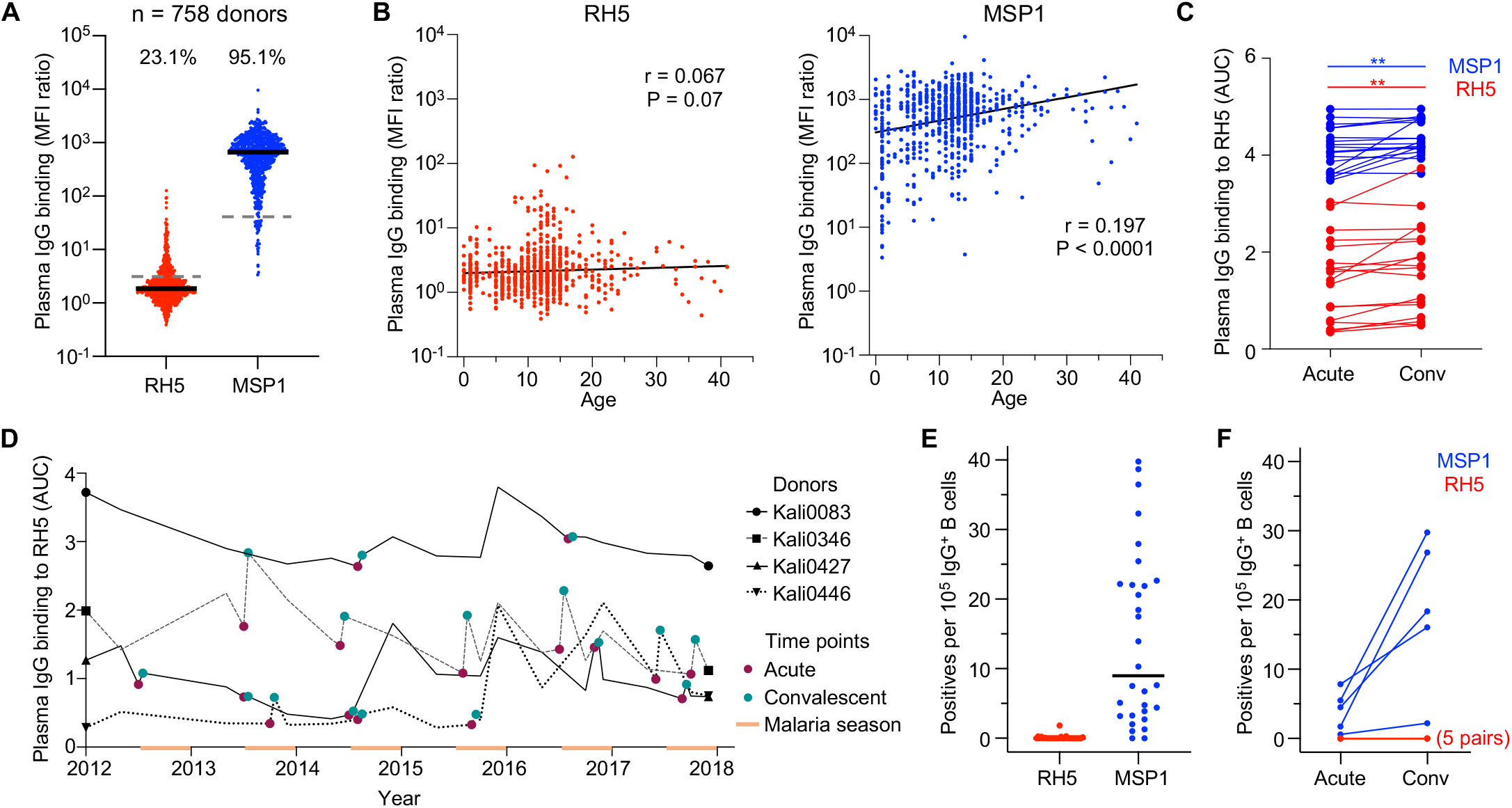
Malaria-exposed individuals have weak B cell responses to RH5 despite repeated infection. A. IgG binding to RH5 and MSP1 of plasma from individuals living in a malaria-endemic region in Kalifabougou, Mali (n = 758 donors). Median fluorescence intensity (MFI) values shown are after division with values for the negative control antigen, CD4. Bars show median values. Dashed lines show the average IgG binding of plasma from 42 US donors. The numbers above each group show the percentage of individuals with plasma binding above this negative control average. B. Cross-sectional analysis of relationship between plasma IgG binding to RH5 and MSP1 with age. Correlation r and P values were determined using Spearman correlation. C. Plasma IgG binding to RH5 of samples from paired acute and convalescent time points. Acute samples were collected upon diagnosis of clinical malaria and convalescent samples were collected ∼1 week after the acute samples. The Wilcoxon sign-rank test was used to analyze changes in binding between the two time points. D. RH5-specific IgG levels in plasma of four Malian donors from whom we isolated RH5-specific mAbs over a period of six years. The orange bars denote the malaria season, which occurs in the latter half of each year. E. Frequency of RH5- or MSP1-positive wells of cultured IgG^+^ B cells from 30 analyzed donors. Two runs where B cells from different donors were pooled into the same wells were excluded from this analysis. Bars show median values. F. Frequency of RH5- or MSP1-positive wells of cultured IgG^+^ B cells from five pairs of acute and convalescent samples. Two pairs are from the same donor (Kali0346), but were taken from different malaria seasons. For RH5, the five pairs are completely overlapping.

Next, we investigated whether RH5-reactive IgG^+^ B cells are detectable in individuals exposed to repeated malaria infection. We selected peripheral blood mononuclear cells (PBMCs) from 30 individuals whose RH5-specific circulating IgG levels were within the top 10% of the Malian cohort. We activated IgG^+^ B cells from these donors polyclonally and screened their supernatants for reactivity to RH5 and MSP1. Out of 10.9 million IgG^+^ B cells in 4,307 wells (2,500 cells/well) from the 30 donors, we identified 1,475 MSP1^+^ wells but only 14 RH5^+^ wells (Figure 1E). To determine if a documented clinical malaria episode triggers a detectable increase in RH5-reactive B cells, we analyzed IgG^+^ B cells from five paired acute and convalescent time points. We detected an increase in MSP1-but not RH5-reactive B cells (Figure 1F). Collectively, these findings suggest that RH5-reactive B cells, while present, are rare in malaria-exposed individuals. Furthermore, merozoite exposure elicits only short-lived RH5-specific IgG antibodies that do not accumulate with repeated malaria infections.

### Malaria exposure elicits rare but potent neutralizing antibodies targeting RH5

Apart from quantifying the antibody response to RH5, we were interested in assessing the quality of the response by isolating RH5-specific mAbs from malaria-exposed individuals. To increase the probability of identifying rare RH5-reactive B cell clones, we used a more labor-intensive but sensitive approach for mAb screening that utilized sequential 384-well oligoclonal and optofluidic monoclonal B cell culture (Dacon et al., 2023). Four individuals with high RH5-specific polyclonal IgG (within the top 10% of the Malian cohort) and multiple PBMC samples available were selected for mAb isolation using this approach. To compare antibodies elicited by natural infection and vaccination, mAbs were also isolated in parallel from 5 malaria-naïve UK adults who had been vaccinated with recombinant RH5 formulated with the adjuvant AS01B (Minassian *et al*., 2021). In these individuals, RH5 vaccination induced long-lasting polyclonal IgG antibodies that only declined slightly 6 months after the final vaccine dose, and we detected >100-fold higher frequencies of RH5-reactive IgG^+^ B cells in vaccinated versus malaria-exposed individuals (Figures 2A and S1A). This difference in B cell frequency was further emphasized by the vastly different number of RH5-specific mAbs we isolated from each group of individuals. For the vaccinees, 164 RH5-specific mAbs were isolated from 0.135 million IgG^+^ B cells whereas, for the malaria-exposed cohort, only 22 RH5-specific mAbs were isolated from 14.4 million IgG^+^ B cells, including B cells from the initial screens described in the previous section (Figure 2B). The mAbs from malaria-exposed donors had acquired more somatic mutations than those from the vaccinated donors, consistent with repeated B cell exposure to RH5 over an extended period of time (Figure 2C). Surprisingly, this did not translate to stronger binding to RH5, based on area under the curve (AUC) measurements of binding to RH5-coated beads (Figure 2D). Upon closer examination, we found that while the number of mutations was associated with improved binding to RH5 in each group, the vaccine-derived mAbs achieved strong binding with fewer somatic mutations than those derived from infection (Figure S1B). We successfully obtained kinetics data on most of the mAbs by surface plasmon resonance (SPR) and found no differences in affinity between the infection- and vaccine-derived mAbs (Figure 2E).

**Figure 2:**
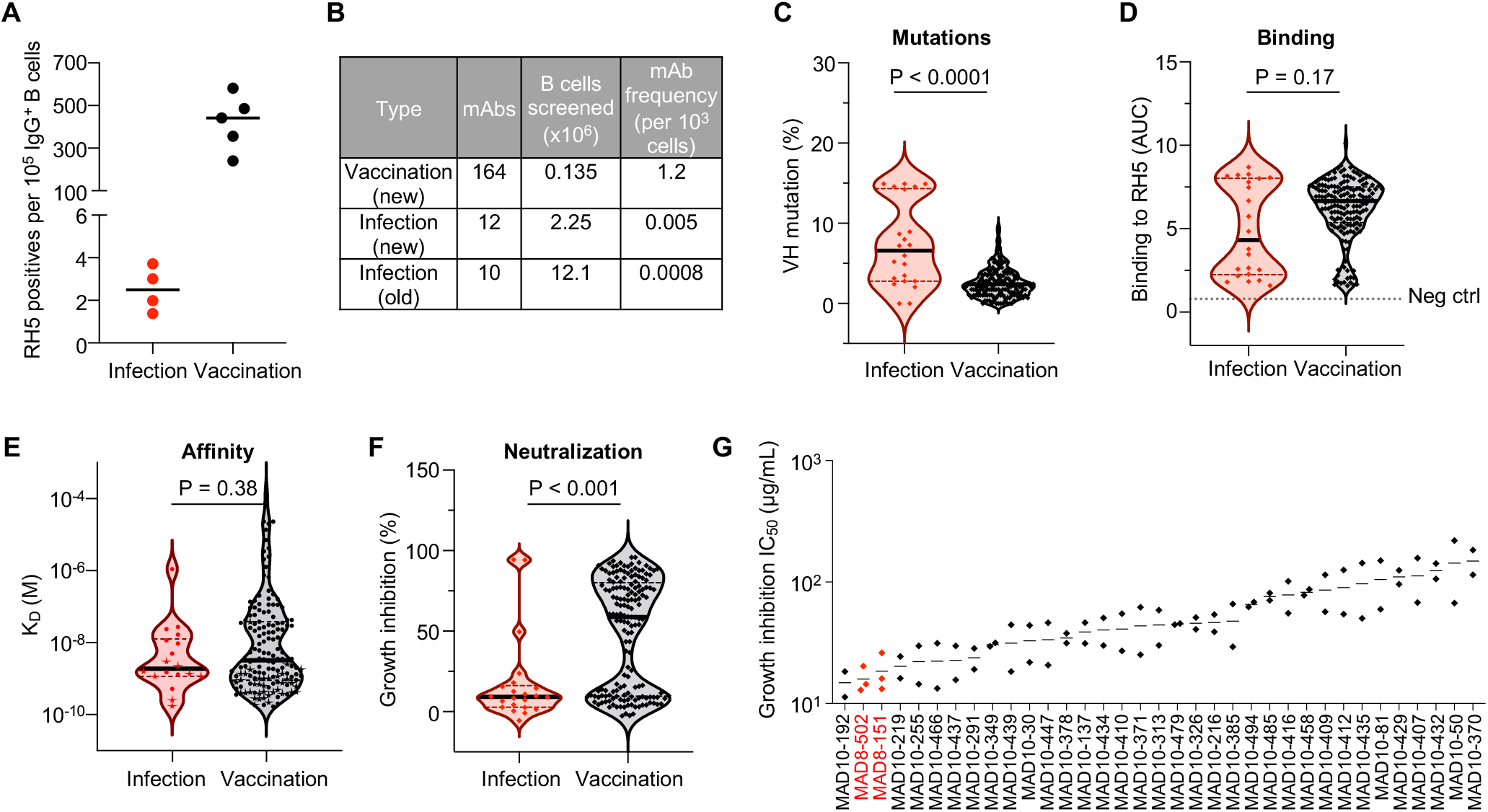
Malaria exposure elicits rare but potent neutralizing antibodies targeting RH5. A. Frequency of RH5-positive wells of cultured IgG^+^ B cells from infected versus vaccinated donors. Bars show median values. B. Number of mAbs isolated from infected versus vaccinated donors. The “new” method refers to the more sensitive but labor-intensive B cell screening approach while the “old” method refers to the original approach used to investigate B cells in Figure 1. C. Heavy chain variable (VH) mutation frequencies of mAbs isolated from infected versus vaccinated donors. Bars shows median values and dashed lines show quartiles. P value was calculated using the Mann-Whitney U test. D. RH5 binding of mAbs isolated from infected versus vaccinated donors. Bars shows median values and dashed lines show quartiles. The dotted line shows the binding of a negative control mAb, VRC01-LS. P value was calculated using the Mann-Whitney U test. E. Binding affinity of mAbs isolated from infected versus vaccinated donors, as measured by the equilibrium dissociation constant (K_D_). Bars show median values and dashed lines show quartiles. P value was calculated using the Mann-Whitney U test. Stars indicate mAbs with a dissociation rate at the limit of detection (1 × 10^-5^ s^-1^), which was the value used to calculate the K_D_ in this graph. Data were obtained for 18/22 infection-derived mAbs and 151/164 vaccination-derived mAbs; the remaining mAbs did not have a sufficient binding signal to the antigen or were acid-denatured during the kinetics workflow. F. Growth inhibition of mAbs isolated from infected versus vaccinated donors. All mAbs were tested at 1 mg/mL. Bars shows median values and dashed lines show quartiles. P value was calculated using the Mann-Whitney U test. G. Growth inhibition half maximal inhibitory concentration (IC_50_) values from titrations of the 35 most potent mAbs. Each point represents an independent experiment. Bars show the mean value.

The panel of 186 RH5-specific mAbs was assessed for function using the growth inhibition assay (GIA), a key *in vitro* reference assay that measures neutralizing activity against blood-stage *P. falciparum* and is highly correlated with *in vivo* protection mediated by RH5 antibodies (Douglas *et al*., 2019; Douglas et al., 2015; Minassian *et al*., 2021). Surprisingly, most of the mAbs from natural infection were non-neutralizing, with only 2 (MAD8-151 and MAD8-502) out of 22 mAbs (9.1%) surpassing the 50% inhibition threshold when tested at 1 mg/mL and a third mAb (MAD8-501) falling just under this threshold (Figure 2F). In contrast, 94 out of 164 vaccine-derived mAbs (57.3%) surpassed this threshold. However, titrations of the 35 mAbs with the highest inhibition values at 1 mg/mL revealed that MAD8-151 and MAD8-502 were among the most potently neutralizing mAbs even when compared to the best vaccination-derived mAbs (Figures 2G and S1C), suggesting that natural infection can elicit potent, albeit uncommon, RH5-specific mAbs.

To determine whether circulating RH5-specific antibodies in infected donors reach sufficient levels to have a detectable functional effect, we screened polyclonal IgG from the source donors of MAD8-151 and MAD8-502, as well as several other Malian donors with high RH5 reactivity, in an antigen-reversal GIA (Miura et al., 2007). In this assay, growth inhibition is compared with and without the adsorption of RH5-specific antibodies to take only RH5-specific antibody activity into account. Only the source donor of MAD8-502, Kali0083, showed a consistent reduction in neutralization when RH5-specific antibodies were adsorbed (Figure S1D), indicating that RH5-specific antibodies rarely show detectable function at the polyclonal level.

### mAbs induced by natural infection primarily target non-neutralizing RH5 epitopes

As the differences in neutralization potency between the vaccination- and infection-derived mAbs could not be explained clearly based on binding strength, we performed an SPR-based epitope binning assay to compare the epitope specificity of mAbs from the two groups. We included reference RH5-specific mAbs with known binding sites to anchor epitope bins to specific regions of RH5 (Alanine *et al*., 2019). Using this approach, we mapped the binding sites of 12/22 infection-derived mAbs and 151/164 vaccine-derived mAbs. We identified 5 major epitope bins, with three bins (I-III) located near the basigin-binding site (i.e., the “top” of RH5) and two bins (IV-V) located at the opposite end of RH5 proximal to the binding site of CyRPA (i.e., the “bottom” of RH5) (Figure 3A).

**Figure 3:**
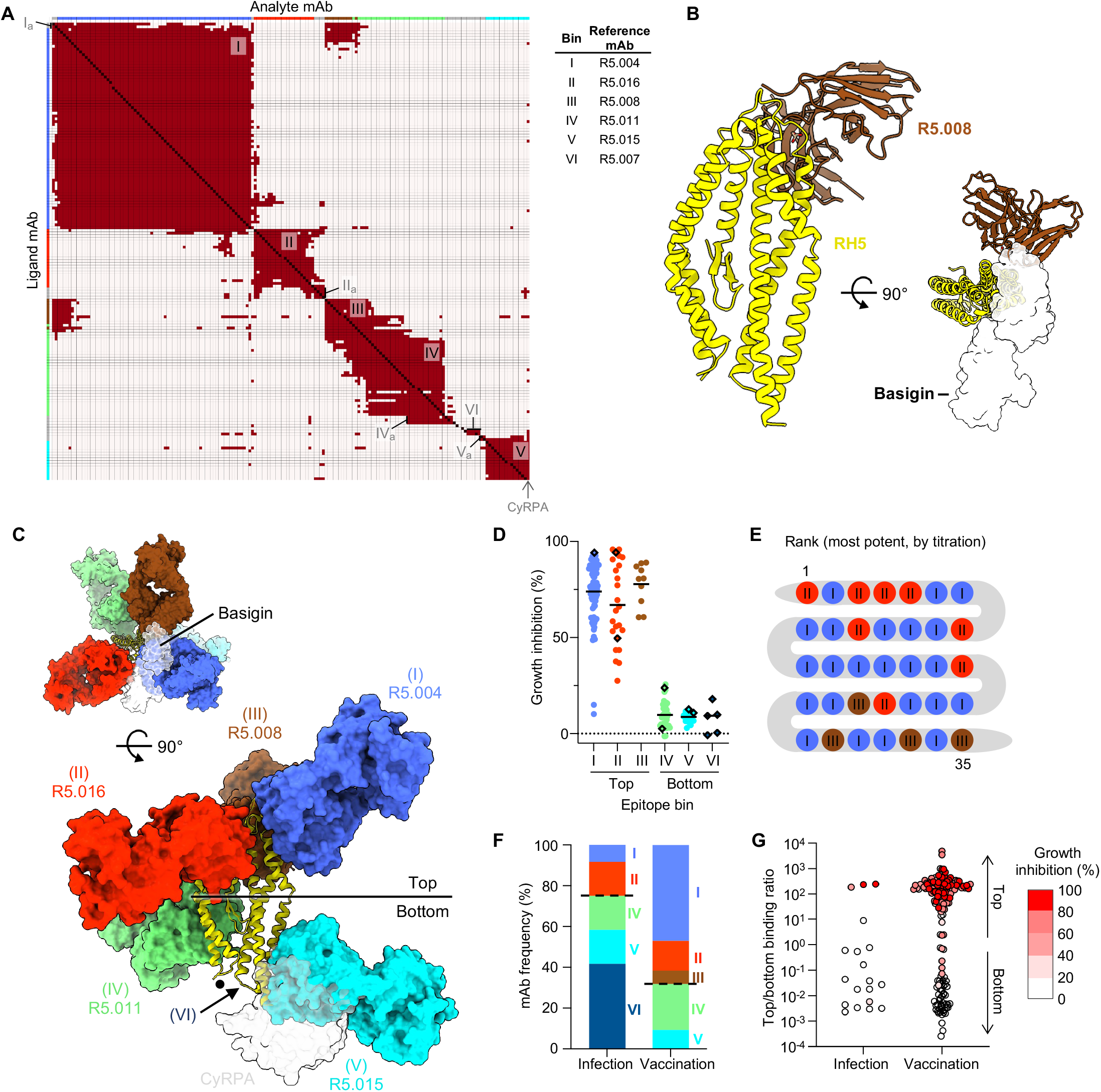
mAbs from natural infection primarily target non-neutralizing RH5 epitopes. A. Heat map showing competition of mAbs targeting RH5. Red squares signify competition between the corresponding analyte and ligand mAb, while cream squares indicate no competition. Adjunct bins are indicated with a subscripted “a”. 12/22 infection-derived mAbs and 151/164 vaccine-derived mAbs were mappable in this assay; the remaining mAbs were incompatible with the workflow, e.g. due to acid sensitivity during the regeneration step. B. Crystal structure of RH5 (yellow) bound to the scFv of R5.008 (brown), each shown in cartoon. R5.008 competes with the binding of basigin (transparent white surface, PDB ID 4U0Q) to RH5. C. Neutralizing and non-neutralizing antibodies target the top and bottom of RH5, respectively. Representative antibody structures for each bin (I = R5.004, PDB ID 6RCU; II = R5.016, PDB ID 6RCV; III = R5.008; IV = R5.011, PDB ID 6RCV; V = R5.015, PDB ID 7PHU) are shown in surface representation on RH5 in cartoon. R5.008 is modelled as a Fab fragment for illustrative purposes. The binding locations of basigin (PDB ID 4U0Q) and CyRPA (PDB ID 8CDD) are shown as transparent surfaces in the top and side views respectively, and the black dot indicates the location of the internal disordered loop of RH5. The approximate location of bin VI is shown by an arrow. D. Growth inhibition of mAbs, subdivided by epitope bin. The adjunct bins were included in the main epitope bins, e.g. bin I_a_ was considered a part of bin I. Bars show mean values. Infection-derived mAbs are shown as diamonds. E. Epitope bins of 35 most potent RH5-specific mAbs, as determined by GIA titration. F. Frequency of infection- and vaccination-derived mAbs in each RH5 epitope bin. The dotted lines separate bins at the top and bottom of RH5. G. mAbs scored by binding proximity to the top of RH5, based on binding to RH5 with bins I-III pre-blocked versus bins IV-VI pre-blocked. Data points are geometric means from two independent experiments. Points are color-coded by GIA score at 1 mg/mL.

All of these major RH5 epitopes have been characterized in detail through crystal structures with reference mAbs (Alanine *et al*., 2019; Ragotte et al., 2022) except bin III, a neutralizing site for which there is a lack of structural information (Alanine *et al*., 2019). We therefore generated an scFv fragment for R5.008 and determined the crystal structure of this mAb bound to RH5 (Figure 3B, Table S1). We find that R5.008 binds to the top half of RH5 to an epitope predominantly formed from the C-terminal two helices (6 and 7) of the RH5-fold, together with the linking loop and the loop between helices 3 and 4. This epitope partly overlaps the binding site for the N-terminal domain of basigin (Wright *et al*., 2014), supporting previous observations that R5.008 directly blocks basigin binding (Alanine *et al*., 2019). Comparison of the R5.008-bound structure with structures of RH5 bound to epitope I (R5.004) and epitope II (R5.016) shows that these three epitopes cover much of the top half of RH5 (Figure 3C). Each of these three epitope bins acts by blocking the binding of RH5 to basigin, either through direct competition as for R5.004 and R5.008 (Figure 3B-C) (Alanine *et al*., 2019), or by blocking the binding of RH5 to basigin-containing membrane protein complexes, as for R5.016 (Jamwal et al., 2022). In contrast, epitope V overlaps directly with the CyRPA-binding site (Ragotte *et al*., 2022), while bin IV is located towards the side at the bottom half of RH5 and contains antibodies which potentiate the effect of neutralizing antibodies (Figure 3C) (Alanine *et al*., 2019). mAbs in a sixth group that were spread over several non-overlapping sites were classified as adjuncts to existing bins (Ia, IIa, IVa or Va) or assigned to a sixth bin (VI) depending on their competition profiles (Figure 3A). Bin VI was mapped to the bottom of RH5 based on competition with CyRPA and the reference mAb R5.007, which targets a disordered loop at the base of the protein (Alanine *et al*., 2019) (Figure 3C).

Strikingly, for both infection- and vaccination-derived mAbs, those targeting the top of RH5 (bins I-III) were predominantly neutralizing, while all mAbs targeting the bottom (bins IV-VI) were non-neutralizing (Figures 3D and S2A-B). Of the 35 most potent mAbs based on the GIA titration (Figure 2G), the top 25 mAbs bound to bins I and II, highlighting these two regions as the most effective RH5 target sites (Figure 3E). Based on the binning analysis, only a minority of infection-derived mAbs (3/12, 25%) targeted neutralizing epitopes at the top of RH5, while the majority of vaccination-derived mAbs targeted these sites (103/151, 68.2%), explaining the discrepancy in potency between the two mAb groups (Figure 3F). To allow the analysis of the epitope specificity of a greater number of infection-derived mAbs, we developed a more sensitive competition assay in which RH5-coated beads were pre-blocked with a cocktail of IgA-switched RH5-specific mAbs targeting the top, bottom, or both regions of RH5 (Figure S2C-E). Binding of each IgG mAb to blocked and unblocked beads was then compared by flow cytometry using a fluorescently labeled anti-IgG secondary antibody. Consistent with the SPR-based analysis, only 4/19 (21.1%) infection-derived mAbs in this expanded panel targeted the top of RH5, while 107/160 (66.9%) of vaccination-derived mAbs targeted this region (Figure 3G). Taken together, these results suggest that natural infection, unlike vaccination, elicits more B cells to non-neutralizing epitopes on the bottom of RH5 away from the basigin-binding site.

### Binding of mAbs targeting bins I and II of RH5 strongly correlates with potency

The functional assessment of a large number of RH5-specific mAbs (n = 186) derived from vaccination and natural infection provided the opportunity to analyze this mAb panel for signatures of potent neutralization. As an initial test, we screened several binding and somatic mutation parameters for correlation with growth inhibition for mAbs from bins I and II, which showed the greatest potency (Figures 3E and 4A). The binding association rate was strongly associated with potency, which is consistent with previous findings on RH5-specific mAbs (Alanine *et al*., 2019) and reflects the need to quickly bind to RH5 to inhibit merozoite invasion, most likely due to transient exposure of RH5 during invasion. The magnitude of binding to RH5-coated beads (as measured by area under the curve, AUC), which captures antibody avidity and durability of the interaction, was most strongly correlated with potency, so we used this feature to analyze the wider mAb panel. Analysis of the mAbs after separation into the major epitope bins (I-V) revealed divergent patterns depending on the bin examined (Figure 4B). Binding to RH5 was strongly correlated with potency for mAbs in bins I and II. The correlation between binding strength and potency was less clear for bin III, but this bin contained relatively few mAbs (n = 10). For mAbs in bins IV and V, increasing binding strength did not increase potency, and even very strong binders in these groups were non-neutralizing. These results suggest that the magnitude of binding to specific epitopes at the top of RH5 are the primary determinants of mAb potency.

**Figure 4:**
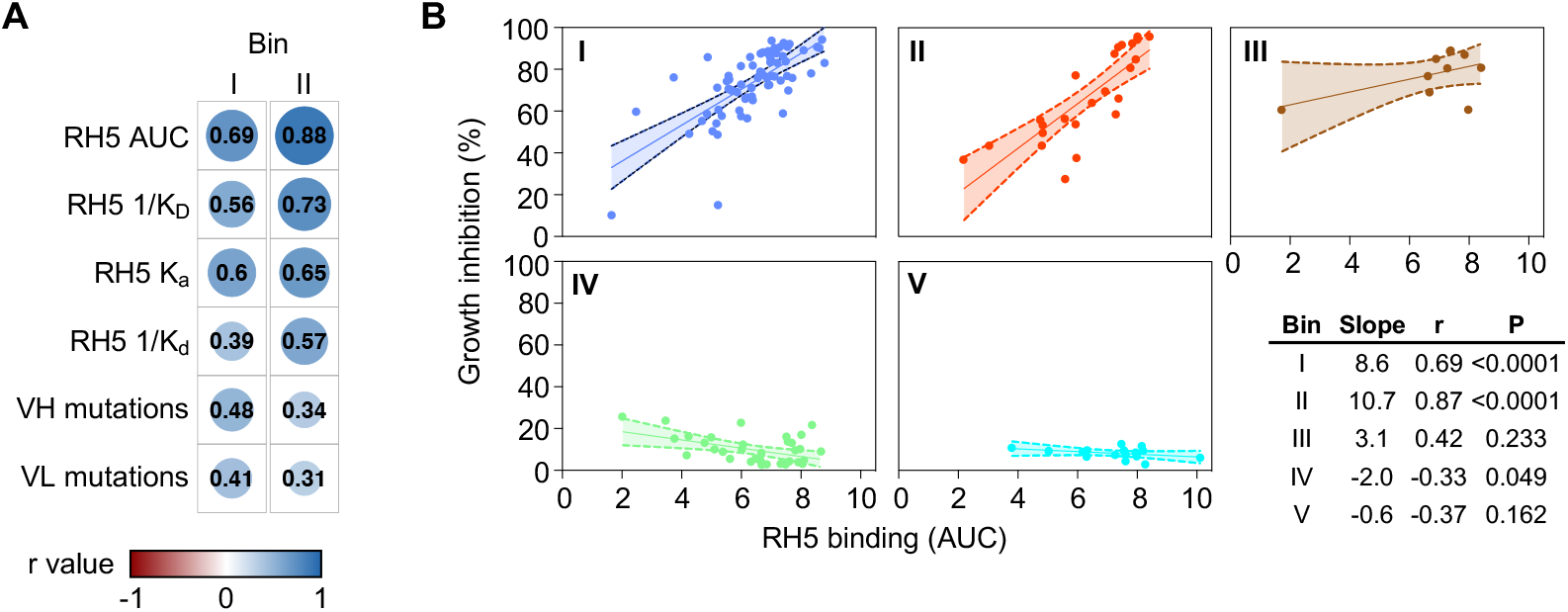
Binding strength of mAbs targeting bins I and II of RH5 is strongly correlated with neutralization potency. A. Spearman correlation between growth inhibition (% at 1 mg/mL) and several binding and sequence parameters for mAbs from bins I and II. The size of the circles and color intensity are proportional to the correlation r value. K_D_, equilibrium dissociation constant; K_a_, association rate constant; K_d_, dissociation rate constant. K_D_ and K_d_ were inverted to allow a direct comparison of positive correlations. Only mAbs from major bins I and II that had measurable values for all 6 parameters were included in this analysis. B. Correlation between growth inhibition and RH5 binding based on AUC for mAbs, subdivided by bin. P and r values were derived from Spearman correlation. Bands show 95% confidence intervals. All mAbs from the major epitope bins (I-V, including adjuncts) with measurable AUC values were considered in this analysis.

### Potent RH5-specific mAbs use diverse VH genes and are clonally dispersed

We compared the genetic backgrounds of mAbs induced by infection and vaccination and the relationship between heavy chain variable (VH) gene usage, target epitope, and potency. Vaccination-derived mAbs using diverse VH genes (VH1-2, VH1-24, VH1-69, VH3-7, VH3-9, VH3-11, VH3-13, VH3-30, VH3-33, VH4-31, VH4-39 and VH4-59) bound the top of RH5 and potently neutralized parasites, indicating a lack of conformational constraints on this target site that necessitate using specific antibody germlines to achieve binding (Figure 5). However, many of the same V genes were also used by mAbs targeting the bottom of RH5. Infection-derived mAbs used substantially fewer VH genes than those derived from vaccination, which was unsurprising given their smaller overall number (Figure 5). These mAbs showed a unique preference for VH3-21, with 9/22 mAbs all targeting the bottom of RH5 using this gene. The bias towards VH3-21 was not solely due to disproportionate expansion of a single clone, as the 9 mAbs originated from 5 distinct clonal lineages and were isolated from 4 different donors (Figure 5). We identified the most expanded lineage in the entire mAb panel here, with 5 mAbs targeting bin VI originating from a single ancestral B cell (Figure S3A). Strikingly, almost every vaccination-derived mAb (162 out of 164 mAbs), including those using common VH genes such as VH4-31 and VH4-39, originated from a distinct B cell ancestor, indicating clonal selection of a wide array of B cells in response to RH5 vaccination.

**Figure 5:**
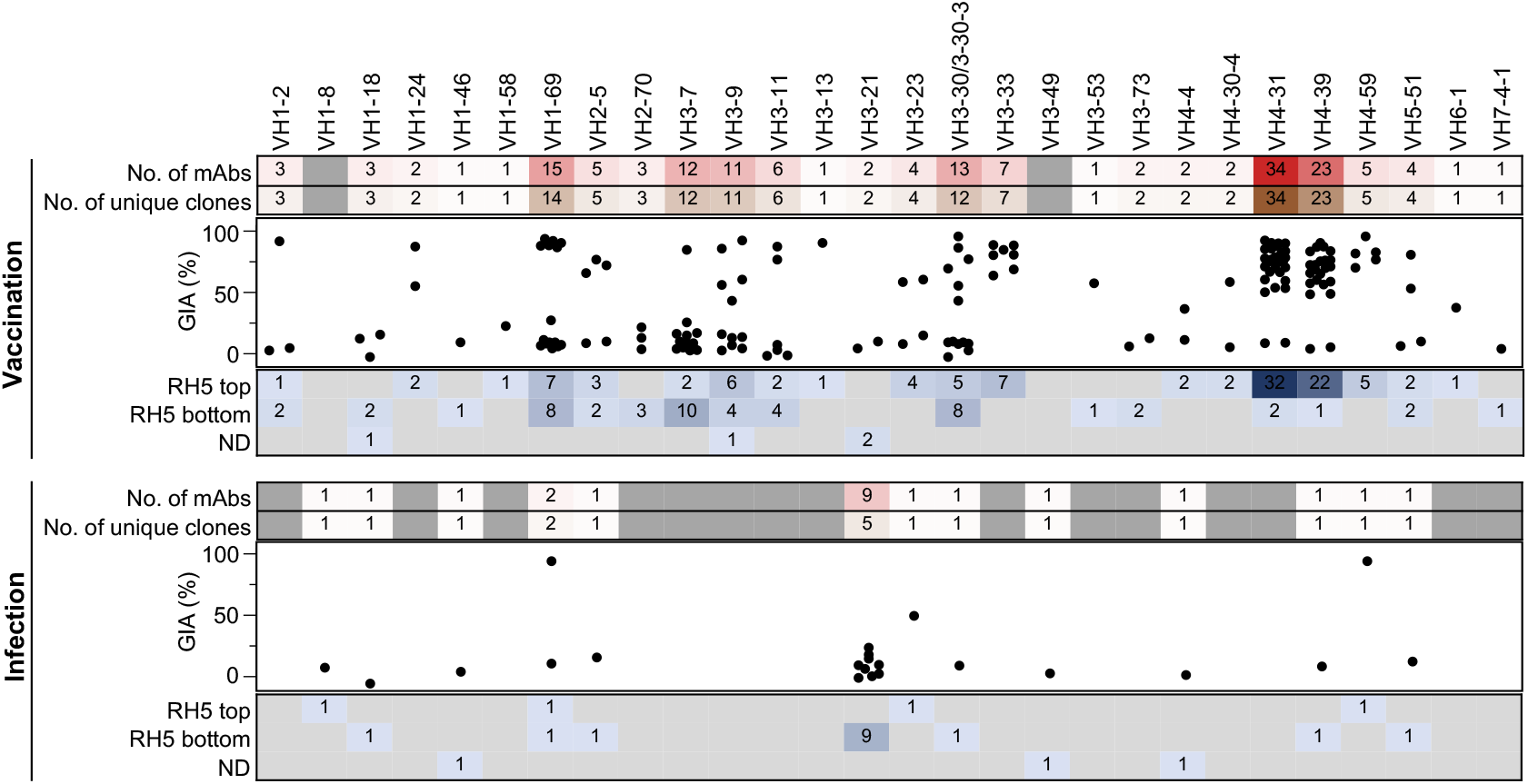
Potent RH5-specific mAbs use diverse VH genes and are clonally dispersed. VH gene usage of mAbs derived from infection and vaccination. Numbers in the bottom panel represent the number of mAbs in each group.

### Identification of a pair of doppelgangers from malaria infection and vaccination

Eleven out of 13 VH genes used by infection-derived mAbs were also used by their vaccination-derived counterparts (Figure 5). Moreover, the two most potent mAbs from natural infection, MAD8-151 and MAD8-502, had similar characteristics to specific vaccine-derived mAbs. MAD8-502 used the same VH gene (VH1-69), targeted the same epitope (bin I), and had similar potency to vaccine-derived MAD10-466 (Figure S3B). Infection-acquired MAD8-151 and vaccination-derived MAD10-255 were even more similar, as they shared all five V(D)J genes, had identical lengths for all six complementarity-determining regions (CDRs), targeted the same region of RH5 (bin II), and had very similar potency (Figure S3B). We therefore determined crystal structures of all four of these antibodies, which allowed us to not only compare them with previously characterized antibodies targeting the same epitopes but also to compare them with one another (Figures 6 and S3, Table S1).

**Figure 6:**
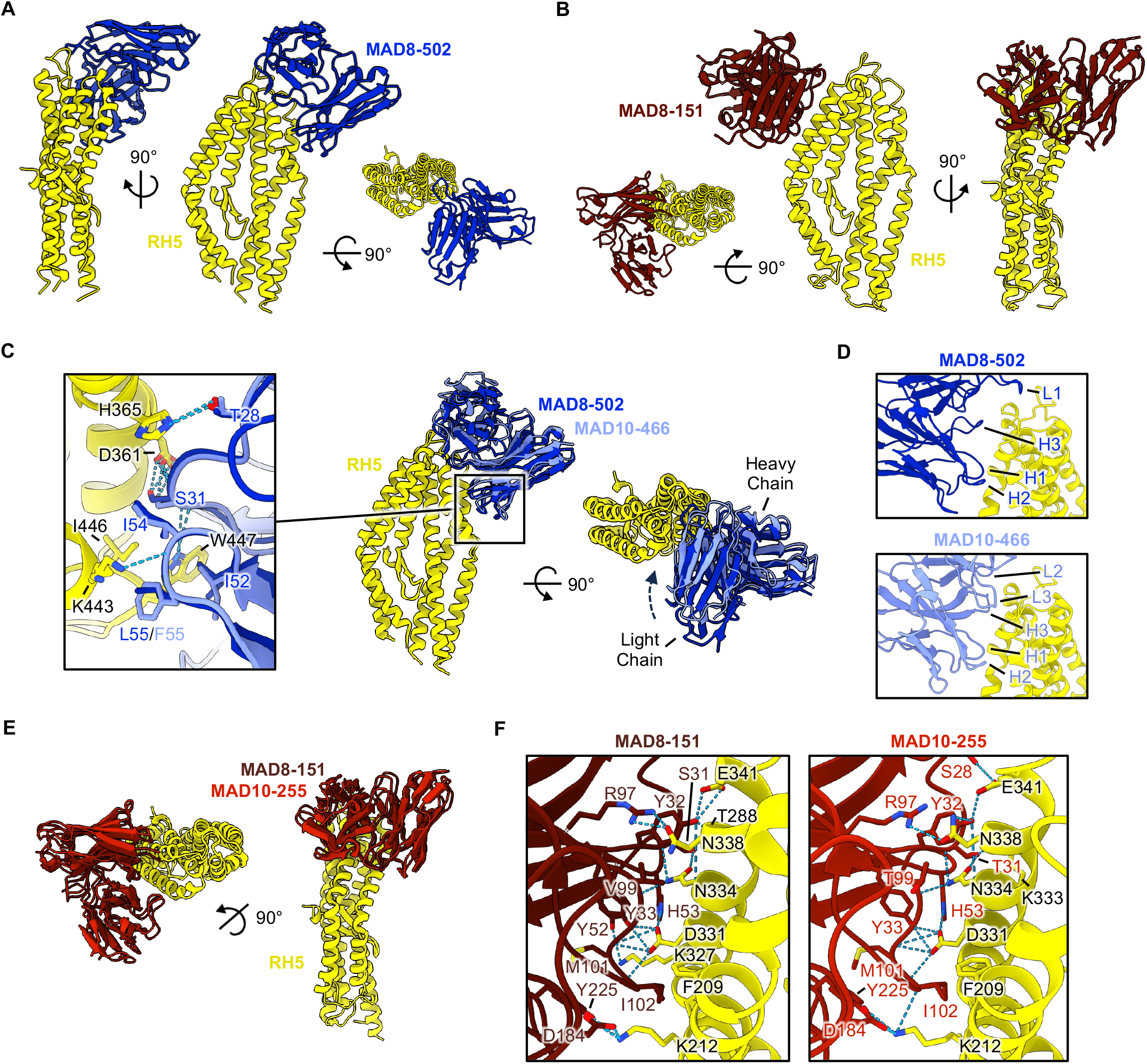
Crystal structures of naturally acquired RH5-targeting mAbs and vaccination-derived counterparts. A. Structure of MAD8-502 scFv (dark blue) bound to RH5 (yellow) from several views, shown in cartoon representation. B. Structure of MAD8-151 scFv (dark red) bound to RH5 (yellow), as in A. C. Overlay of naturally-acquired MAD8-502 (dark blue) and vaccine-derived MAD10-466 (light blue) bound to RH5 (yellow) illustrating similar binding modes. Only the variable domain of the MAD10-466 Fab fragment is shown for simplification. Shared intermolecular contacts made by CDRs H1 and H2 of each mAb are shown in the expanded panel inset, with residues shown as sticks and hydrogen bonds with dashed lines. D. Comparison of the binding interfaces between MAD8-502 and MAD10-466 to RH5 demonstrating shared use of CDRs H1 and H2 to interact with RH5 but variable use of CDRs H3, L1, L2 and L3. E. Overlay of doppelgangers MAD8-151 scFv (naturally-acquired, dark red) and MAD10-255 (vaccination-derived, red) bound to RH5 (yellow). F. View of the almost identical binding interface between MAD8-151 and MAD10-255 with RH5. Residues taking part in the binding interface are labelled and shown as sticks while hydrogen bonds are shown as dashed lines. In the MAD8-151 panel, RH5 residues are numbered as in wild-type RH5 to allow for comparison to MAD10-255, while the deposited structure uses numbering of the RH5ΔNL construct (Table S2).

To enable crystallization, we prepared single-chain variable fragments (scFv) for the naturally acquired antibodies MAD8-502 and MAD8-151 and crystallized complexes of each scFv bound to RH5 (Figures 6A-B). As expected from epitope binning, MAD8-502 bound to a site on the top half of RH5, with an epitope consisting of helices 4 and 6, and the loops joining helices 3-4 and 6-7 (Figure 6A). This site overlaps with the binding site for R5.004, with MAD8-502 binding to RH5 with a very similar angle of approach (Figure S3C), and sharing a large number of contact residues (Table S2) (Alanine *et al*., 2019). In contrast, MAD8-151 bound to an epitope on the opposite face of the top of RH5, formed from the anti-parallel helices 2 and 3 (Figure 6B). This overlaps with the epitope of R5.016 (Table S2) (Alanine *et al*., 2019). However, MAD8-151 binds to RH5 with an angle more like that adopted by the mouse mAb 9AD4 (Wright *et al*., 2014) than R5.016 (Figure S3D) and makes more contacts with RH5 helix 3 than R5.016, which has a preference for helix 2.

To enable a comparison with MAD8-502 and MAD8-151, we next determined the structures of vaccine-derived antibodies MAD10-466 and MAD10-255 bound to RH5 using fragment antigen-binding (Fab) or scFv fragments respectively to enable crystallization (Figures S3E and S3F). Aligning the structures of these vaccine-derived antibodies to their naturally acquired counterparts reveals that each pair bind to RH5 with very similar poses (Figures 6C and 6E). In the case of the MAD8-502 and MAD10-466 pair, binding similarity is predominantly within their heavy chains, where interactions made by CDRs H1 and H2 to RH5 are conserved (Figure 6C, Table S2). These CDRs are encoded by the VH1-69 gene, which the mAbs share, and utilize a conserved hydrophobic patch in CDR H2 (residues I52, I54 and F/L55) to bind RH5. While CDRs H3 in both antibodies make several hydrophobic and hydrogen bond interactions with RH5, and share an interaction with R357 of RH5, these occur due to different residues, reflecting differences in gene usage and amino acid sequences despite their CDRs being similar lengths. Indeed, MAD10-466 binds more intimately with RH5 than MAD8-502, with the light chain variable domain rotated slightly towards RH5 (Figure 6C), enabling CDRs L2 and L3 to make contacts (Figure 6D). Meanwhile, only CDR L1 from the light chain of MAD8-502 interacts with RH5 (Figure 6D, Table S2).

The naturally-acquired MAD8-151 and vaccine-derived MAD10-255 (bin II), which both target the most potent epitope of RH5 (Alanine *et al*., 2019), are remarkably similar, sharing all five V(D)J genes (Figure S3B). An overlay of RH5 bound to MAD8-151 and MAD10-255 shows that these two antibodies bind with an almost indistinguishable binding mode, overlapping with a backbone root mean square deviation (RMSD) of 0.57 Å (Figure 6E). Moreover, contact residues between each antibody and RH5 are almost entirely conserved, using the same residues from CDRs H3 and L2-3 (Table S2). Both antibodies make identical hydrogen-bond mediated binding interactions with D331 of RH5, and several nearly identical interactions with residues F209, K212, N334, N338, and E341 of RH5 (Figures 6F, Table S2). We therefore dubbed MAD8-151 and MAD10-255 “doppelgangers” since they have similar structural and functional properties, but distinct geographical and biological origins. These findings indicate that, at least in these two cases, natural malaria infection is capable of eliciting B cells with similar characteristics to those induced by vaccination with recombinant RH5, despite differences in the mode of antigen exposure and donor origin.

## DISCUSSION

A key immune evasion strategy employed by pathogens is the concealment of conserved functional sites through masking by other molecules, protein folding, or antigen sequestration in intracellular organelles. Here, we find that *P. falciparum* blood-stage merozoites successfully avoid triggering a strong neutralizing antibody response to a conserved and functionally important antigen, RH5, despite frequent infection of the same individuals over multiple years. Based on the low frequency of RH5-reactive memory B cells and long-lived antibodies, exposure to RH5 during *P. falciparum* infection appears unable to consistently induce robust germinal center responses, although extensive somatic mutations in rare RH5-specific mAbs suggests that germinal center maturation does occur on limited occasions. These findings are consistent with previous work reporting low titers of circulating RH5-specific antibodies in individuals living in malaria-endemic regions (Douglas *et al*., 2011; Partey *et al*., 2018; Tran *et al*., 2014; Willcox *et al*., 2021). Additionally, this study identifies a second obstacle to generating a protective anti-RH5 antibody response: the antibodies that overcame the first hurdle and were generated in response to infection, including the only clonally expanded family we isolated, were more commonly found to target non-neutralizing “bottom” regions of RH5 distal from the basigin-binding site. This observation aligns with previous data suggesting that CyRPA and RIPR detach from RH5 through disassembly of the CyRPA-RH5 interaction upon binding of the RH5 complex to red blood cells (Wong *et al*., 2019), potentially leaving RH5 attached to basigin with only the bottom of this protein exposed to circulating B cells. While a class of mAbs (IV) that binds to an epitope at the side of RH5 towards the bottom half of the protein can potentiate the neutralizing activity of antibodies targeting the top of the protein (Alanine *et al*., 2019), only mAbs that predominantly target epitopes at the top of the protein that interfere with binding to the basigin complex (Jamwal *et al*., 2022) have direct and consistent neutralizing activity. As current RH5 vaccines include both top and bottom regions of the protein and can activate B cells targeting both sites, this observation supports the investigation of next-generation RH5 vaccines that consist of only the top of the protein or mask bottom epitopes to avoid activating non-neutralizing B cells in individuals with prior malaria exposure. The identification of rare B cells from infected donors that target neutralizing sites on the top of RH5, as well as shared V gene usage between infection-and vaccination-derived mAbs, provides encouragement that malaria-exposed individuals contain a suitable B cell repertoire for activation by such a vaccine.

The analysis of a large panel of mAbs derived from RH5 vaccination and natural infection allowed us to identify precise rules governing the potency of RH5-specific antibodies. These findings build upon previous work on a smaller number of mAbs from vaccinated humans or animals, which indicated that antibodies blocking interactions with the basigin receptor complex can potently inhibit merozoite invasion (Alanine *et al*., 2019; Douglas et al., 2014; Jamwal *et al*., 2022). Here, we identified two regions of RH5, denoted as bins I and II, as the target sites of the 25 most potent mAbs out of a panel of 186 mAbs, with a third neutralizing epitope, structurally characterized here for the first time (bin III), acting as the target of a smaller number of mAbs. Moreover, we found a strong correlation between binding and potency for mAbs within bins I and II. These findings identify a conceptually simple goal for a next-generation RH5 vaccine: to elicit high-affinity antibodies against these two sites. Furthermore, they also provide tools to screen and down-select new vaccine candidates based on this singular feature. In terms of gene usage, a variety of germline VH genes were shared by neutralizing and non-neutralizing mAbs, suggesting that triggering specific germlines may be less beneficial than an epitope-driven approach for designing a next-generation RH5 vaccine. The activation of almost entirely clonally dispersed B cells targeting neutralizing epitopes on RH5 bodes well for the goal of developing an RH5 vaccine that is effective in individuals with different genetic backgrounds, as it suggests that the vaccine does not need to selectively trigger rare clones with specific features in order to elicit a successful antibody response.

### Limitations of the study

A limitation of this study is the relatively few RH5-specific mAbs isolated from malaria-exposed individuals. Despite screening a large number of B cells (∼14 million) from 30 donors that were selected out of a large cohort (n = 758) based on high circulating levels of polyclonal RH5-specific antibodies, we were only able to isolate 22 distinct RH5-specific mAbs. The main constraint here was the low frequency of detectable RH5-reactive B cells induced by natural infection. It would be ideal to validate the ratio of RH5-reactive B cells from natural infection targeting the top and bottom of the protein with a larger number of mAbs, but we anticipate that this will require a substantial, dedicated effort to screen even more samples from malaria-exposed donors.

## ACKNOWLEDGMENTS

We thank the blood sample donors from the Kalifabougou cohort and VAC063 trial. We thank Ludmila Krymskaya for sorting of PBMC samples. This work was supported by the Division of Intramural Research and the Vaccine Research Center, National Institute of Allergy and Infectious Diseases (NIAID), National Institutes of Health (NIH) (LTW, AJRC, KM, AD, LP, CT, SL, RAS, PDC, CAL, JT). The VAC063 clinical trial, monoclonal antibody isolation and R5.008 structure determination were made possible IN part through support provided by the Infectious Disease Division, Bureau for Global Health, United Agency for International Development (USAID), under the terms of the Malaria Vaccine Development Program (MVDP) (AID-OAA-C-15-00071 and 7200AA20C00017). For the VAC063 clinical trial, Leidos Inc. was the prime contractor; some of the structural work was supported by USAID under the terms of GH-BAA-2018-Addendum03. MKH and BF are funded by a Wellcome Investigator Award (220797/Z/20/Z). This work was funded in part by the Wellcome Trust (grant 206194). The GIA activity was supported by an Interagency Agreement (AID-GH-T-15-00001) with the USAID MVDP. CMN held a Sir Henry Wellcome Postdoctoral Fellowship [209200/Z/17/Z], and SJD is a Jenner Investigator and held a Wellcome Trust Senior Fellowship [106917/Z/15/Z]. The findings and conclusions are those of the authors and do not necessarily represent the official position of USAID.

## AUTHOR CONTRIBUTIONS

Conceptualization: JT, LTW, AJRC, MKH, BF. Methodology: LTW, AJRC, BF, KM, CWP, MKH, JT. Formal analysis: LTW, AJRC, BF, KM, AD, CT, CWP, MKH, JT. Data curation: SL, SD, DD, KK, AO. Investigation: LTW, AJRC, BF, KM, AD, NMS, CC, LP, MM, JRB, KM, CT, CWP, CMN, SES. Resources: DCN, FEHL, AMM, PDC, SJD. Writing – original draft: LTW, AJRC, BF, MKH, JT. Writing - review & editing: all authors. Visualization: LTW, AJRC, BF, KM, MKH, JT. Supervision: FEHL, DEG, BT, RAS, PDC, GJW, SJD, CAL, MKH, JT. Funding acquisition: PDC, GJW, SJD, CAL, MKH, JT.

## MATERIALS AND METHODS

### Human clinical specimens

Clinical specimens were sourced from a longitudinal study of 758 malaria-exposed subjects aged 1 month to 41 years (49.9% female) in the rural village of Kalifabougou, Mali (Tan et al., 2021; Tran *et al*., 2014) or the VAC063 study (a multi-center, non-randomized, open-label, dose escalation Phase I/IIa clinical trial evaluating the safety, immunogenicity, and efficacy of RH5.1 formulated with the adjuvant AS01B in 88 healthy, malaria-naïve subjects 18-45 years of age, 62.5% female, in the United Kingdom; ClinicalTrials.gov: NCT02927145) (Minassian et al., 2021). For the Malian subjects, PBMCs and plasma were isolated from venous blood at the following timepoints: before and after each malaria season, when a malaria infection was diagnosed via referral or biweekly or monthly scheduled visits (acute), and 7-10 days after the confirmed malaria infection (convalescent). PBMCs from all timepoints were used to isolate mAbs. For the VAC063 subjects, mAbs were isolated from PBMCs collected 12 weeks after the final immunization in Groups 1 and 4 (which respectively received three monthly vaccinations of 2 and 50 µg of RH5.1/AS01B).

### Ethics approval

The Kalifabougou cohort study was approved by the Ethics Committee of the Faculty of Medicine, Pharmacy and Dentistry at the University of Sciences, Technique and Technology of Bamako, and the Institutional Review Board of the National Institute of Allergy and Infectious Diseases, National Institutes of Health (NIH IRB protocol number: 11IN126; https://clinicaltrials.gov/; trial number NCT01322581). Written informed consent was obtained from participants or parents or guardians of participating children before inclusion in the study. The RH5 vaccination study was approved by the UK NHS Research Ethics Service (Oxfordshire Research Ethics Committee A, Ref 16/SC/0345), as well as the UK Medicines and Healthcare products Regulatory Agency (Ref 21584/0362/001-0011).

### Production of recombinant malaria antigens

*P. falciparum* RH5 used for antibody isolation was expressed as a soluble recombinant protein containing a C-terminal rat Cd4 domains 3+4 tag followed by a site for BirA enzymatic biotinylation and a 6-histidine tag for purification essentially as described (Galaway et al., 2019). The *P. falciparum* merozoite antigen MSP-1 was produced in a similar manner (Crosnier et al., 2013). A protein consisting of only rat Cd4 domains 3+4 was used as a control. Proteins were enzymatically biotinylated during expression by co-transfection with plasmids encoding a secreted *Escherichia coli* BirA construct (Elton et al., 2019). All proteins were expressed as secreted proteins by transient transfection in the human HEK293E cell line grown in suspension as described (Zenonos et al., 2014). His-tagged proteins were purified from transient transfection supernatants on HisTrap HP columns (GE Healthcare) using an ÄKTAxpress (GE Healthcare) according to the manufacturer’s instructions. RH5 for kinetics and epitope binning experiments was produced as previously described (Ragotte *et al*., 2022) using a stably transfected *Drosophila* S2 cell line and purified by C-tag affinity purification and size exclusion chromatography. RH5ΔNL (a construct of RH5 encompassing residues K140-K247 and N297-Q526, thereby lacking its flexible N-terminus and internal loop, with substitutions T216A and T299A [to remove potential glycosylation sites] and C203Y [of the 7G8 *Plasmodium falciparum* strain], and a C-terminal His-tag) was expressed and secreted from a stable S2 cell line (ExpreS^2^ion Biotechnologies) in EX-CELL® 420 Serum Free Medium (Sigma Aldrich) (Wright *et al*., 2014). After 3-4 days, the culture supernatant was harvested and adjusted to pH 8 with Tris, spun at 9,000 g for 15 mins and 0.45 μm filtered, then incubated with Ni Sepharose™ excel resin (Cytiva) for 2 hours. Beads were washed with 5 column volumes of TBS (20 mM Tris pH 8.0, 150 mM NaCl), and 20 column volumes of wash buffer (20 mM Tris pH 8, 500 mM NaCl, 20 mM imidazole), then bound proteins eluted with elution buffer (20 mM Tris pH 8, 150 mM NaCl, 500 mM imidazole). Eluted proteins were diluted 1:1 in ConA binding buffer (20 mM Tris pH 7.5, 500 mM NaCl, 1 mM MnCl2, 1 mM CaCl2), then incubated with ConA Sepharose 4B resin (Cytiva) overnight at 4°C, after which the unbound fraction containing RH5ΔNL was recovered. RH5ΔNL was further purified by gel filtration using an Superdex 200 Increase 10/300 column into SEC buffer (20 mM HEPES pH 7.5, 150 mM NaCl, 5% glycerol).

### Plasma and mAb binding to antigen-coated beads

Streptavidin beads (Spherotech) were incubated with biotinylated RH5 (50 µg/mL) and MSP1 (10 µg/mL) for 30 minutes at room temperature, washed with PBS + 0.5% BSA, blocked with CD4 (10 µg/ml; negative control antigen) for 30 minutes at room temperature, washed, mixed, aliquoted, and frozen at -80°C. To measure plasma binding, 1:20 diluted Malian plasma was incubated with thawed antigen-coated beads for 30 minutes at room temperature, washed, and stained with 2.5 µg/mL goat anti-human IgG Alexa Fluor-647 (AF647) secondary antibody (Jackson Immunoresearch). To measure mAb binding, eight mAb dilutions (25, 3.57, 0.51 µg/mL, etc.; 7-fold serial dilutions) were incubated with antigen-coated beads for 30 minutes at room temperature, washed, and incubated with IgG-AF647. After a final wash, plasma- or mAb-bound beads were read with the iQue Screener Plus (Intellicyt) high-throughput flow cytometer and data were analyzed with FlowJo (Tree Star). Malian plasma data were normalized by dividing RH5 or MSP1 median fluorescence intensity (MFI) by CD4 MFI. For mAb titrations, AUC values were calculated using GraphPad Prism and normalized based on interexperimental variance of positive control RH5-specific mAb R5.016 (Alanine *et al*., 2019) with the first experiment’s R5.016 AUC values for RH5 binding being used as the anchors by which all subsequent R5.016 values were divided; anti-HIV-1 mAb VRC01 (Wu et al., 2010) was included as a negative control.

### Isolation of RH5-specific B cells

RH5-specific memory B cells (MBCs) and plasmablasts (PBs) were identified using the Beacon Optofluidic System as previously described (Cho et al., 2021). Briefly, cryopreserved PBMCs were thawed and stained for 20 minutes at 4°C in PBS with LIVE/DEAD Fixable Aqua (Thermo Fisher Scientific) prior to staining for 20 minutes at 4°C in PBS + 1% FBS with the following panel: CD3-BV510, CD14-BV510, CD27-AF488, CD38-APC/Cy7, CD56-BV510 (BioLegend), CD19-ECD (Beckman Coulter), IgA-AF647 (Jackson Immunoresearch), IgM-PerCP-Cy5.5, and IgD-PECy7 (BD Biosciences). Sorted single CD3^-^CD14^-^CD56^-^CD19^+^IgD^-^IgM^-^IgA^-^IgG^+^ MBCs were cultured via two methods: 1) 2,500 MBCs/well cultured in a proprietary cytokine cocktail (Berkeley Lights) for 6 days in 96-well U-bottom plates or, 2) 100-250 MBCs/well and 10,000 3T3-CD40L feeder cells/well (Huang et al., 2013; Moir et al., 1999) cultured in I10 media (Iscove’s modified Dulbecco’s Medium, 10% FBS, 1:1000 MycoZap; Thermo Fisher and Lonza) supplemented with 100 ng/mL IL-21 (Gibco) and 0.5 µg/mL R848 (Mabtech) for 10 days in 384-well plates. After stimulation, culture supernatants were screened for binding to antigen-coated beads as described above. MBCs from positive wells were suspended in either Plasma Cell Survival Media (PCSM, Berkeley Lights) or I10 media supplemented with IL-21 and R848, loaded onto an OptoSelect 11k chip, and individually moved into nanoliter-volume pens using opto-electropositioning (OEP) light cages drawn by the Beacon’s Cell Analysis Suite (Berkeley Lights). Channels were then flooded with 7 µm streptavidin beads (Spherotech, SVP-60-5) coated with 50 μg/mL of biotinylated RH5 and suspended in a cocktail of 2.5 μg/mL IgG-AF647. Over a 30-minute time course, B cells producing RH5-specific antibodies were detected by the appearance of fluorescent ‘blooms’. Live singlet CD3^-^CD14^-^CD56^-^CD19^+^CD27^+^CD38^+^ PBs were sorted into PCSM and screened in a similar manner as above in PCSM. Positive B cells were individually exported using OEP light cages into 96-well plates containing lysis buffer and frozen at −80°C.

### mAb sequence analysis and production

Antibody heavy (VH) and light (VL) chains were PCR-amplified and sequenced as previously described (Cho et al., 2021; Wang et al., 2020). Sequence analysis, including determination of VH/VL genes and percentage of somatic mutations, was performed using the International Immunogenetics Information System (IMGT) database. Clonal analysis of the mAb panel was performed with SHazaM and Spectral Clustering for clOne Partitioning (SCOPer) from the Immcantation analysis framework (Nouri and Kleinstein, 2020). Briefly, the VH sequences were analyzed by SHazaM to determine the threshold for clonal grouping (using the gmm model), which was used by SCOPer to determine the clonal relationship between the mAbs. Only MAD8-652 could not be analyzed in this system, but this mAb was unambiguously identified as belonging to a unique clone as it used a different VH gene and CDRH3 length from all other mAbs isolated from its source donor. Antibody VH or VL sequences were cloned into plasmids containing an IgG1 or relevant light chain backbone (GenScript) and transfected into Expi293 cells (Thermo Fisher Scientific). Recombinant IgG was purified using HiTrap Protein A columns (GE Healthcare Life Sciences). To produce antibody Fabs, heavy chain plasmids encoding only the VH and CH1 (domain 1 of the constant region of the immunoglobulin heavy chain) were synthesized and used to transfect Expi293 cells along with light chain plasmids. Fab fragments were purified by anti-CH1 resin (Thermo Fisher Scientific). scFv fragments for MAD8-151, MAD8-502 and MAD10-255, consisting of VH and VL domains joined by a flexible 15-residue long glycine-rich linker sequence, were transiently expressed in Expi293 cells and purified by His-tag affinity columns. An scFv construct of R5.008 (Alanine *et al*., 2019) comprising the variable heavy chain and light chains joined with the 15-residue linker was transiently expressed in a secreted form using FreeStyle™ 293-F cells (Thermo Fisher) in FreeStyle™ F17 Expression Medium supplemented with L-glutamine and 1x MEM non-essential amino acids (Gibco). After 6 days, cultures were purified by His-tag affinity as performed for the other scFv fragments. To isolate monomeric scFv, eluted proteins were further purified by gel filtration on an S200 Increase 10/300 into 20 mM HEPES pH 7.5, 150 mM NaCl at room temperature.

### HLA typing of donor cDNA

When B cells from multiple donors were pooled for optofluidic screening, HLA-typing was required to match isolated mAbs to donor source. HLA-typing was carried out as described previously (Dacon *et al*., 2023). Briefly, the ScisGo^®^-HLA-v6 kit (Scisco Genetics Inc., Seattle WA) was used for amplicon-based sequencing of cDNA from individual cell isolates. This entailed uses a two-stage PCR for locus amplification and sample barcoding. Resulting reactions were then purified and subjected to MiSeq using Illumina Version 2 chemistry with 500-cycle, paired-end sequencing (Illumina, San Diego, CA). Data were assembled with Sciscloud^®^ software (Scisco Genetics Inc., Seattle WA),

### Antibody kinetics

All kinetics experiments were performed with the Carterra LSA. A CMDP chip (Carterra 4282) was primed with filtered and degassed 25 mM MES buffer. The chip was conditioned with 50 mM NaOH, 500 mM NaCl and 10 mM Glycine (pH 2.0) (Carterra 3640), and activated with a mixture of 400 mM 1-ethyl-3-(3-dimethylaminopropyl) carbodiimide hydrochloride and 100 mM N-hydroxysuccinimide (Thermo Fisher Scientific). The activated chip was lawned with goat anti-human IgG Fc (50 µg/mL) (Jackson ImmunoResearch) in 10 mM sodium acetate (pH 4.5) (Carterra 3802) with 0.05% Tween (Carterra 3631) before blocking with 1 M ethanolamine (pH 8.5) (Carterra 3626) and washing with 10 mM Glycine (pH 2.0). Next, mAbs were diluted to 100 ng/mL in HEPES-buffered saline Tween-EDTA (HBSTE) with 0.5mg/mL BSA. This array was printed onto the chip by capture. RH5 was then injected onto bound mAbs in eight increasing concentrations, up to a top concentration of 500 nM, following a 10-minute association and 30-minute dissociation protocol. The SPR results were exported to Kinetics Software (Carterra) and analyzed as nonregenerative kinetics data to calculate association rate constant (Ka), dissociation rate constant (Kd), and equilibrium dissociation constant (KD) values.

### Epitope binning

Epitope binning experiments were performed with the Carterra LSA. A HC30M chip (Carterra 4279) was primed with filtered and degassed 25 mM MES buffer. The chip was conditioned with 50 mM NaOH, 500 mM NaCl and 10 mM Glycine (pH 2.0) (Carterra 3640), and activated with a mixture of 400 mM 1-ethyl-3-(3-dimethylaminopropyl) carbodiimide hydrochloride and 100 mM N-hydroxysuccinimide (Thermo Fisher Scientific). The activated chip was coupled directly to mAbs diluted to 10 µg/mL in 10 mM sodium acetate (pH 4.5) (Carterra 3802) with 0.05% Tween (Carterra 3631), before blocking with 1 M ethanolamine (pH 8.5) (Carterra 3626) and washing with 10 mM Glycine (pH 2.0). The coupled antibody array was then subjected to sequential injections of 50 nM RH5 and 10 µg/ml sandwiching antibody, both diluted to final concentration in HEPES-buffered saline Tween-EDTA (HBSTE) with 0.5 mg/mL BSA. The protocol followed involved 5 minutes for antigen injection, 5 minutes for monoclonal antibody injection and 1 minute for dissociation, before glycine-mediated chip regeneration, allowing for the injection of the next antibody in the array. The SPR results were exported to Epitope Software (Carterra) to identify competing or sandwiching pairs of monoclonal antibodies.

### Epitope localization assay

Streptavidin beads (Spherotech) were incubated with biotinylated RH5 (50 µg/mL) for 30 minutes at room temperature, washed with PBS + 0.5% BSA, blocked with CD4 (10 µg/ml; negative control antigen) for 30 minutes at room temperature, washed, and subsequently incubated with high-affinity anti-RH5 mAbs expressed in the IgA1 isotype. Several bead populations with differing intrinsic fluorescent intensities were used to set up this multiplex assay. One set of beads was exclusively incubated with top-binding anti-RH5 mAbs, another with only bottom-binding anti-RH5 mAbs, and yet another with both top and bottom-binding anti-RH5 mAbs. A fourth set of beads was left unblocked. After 30 minutes incubation, bead batches were washed and mixed. All anti-RH5 mAbs, in IgG format, were incubated with these bead mixes for 30 minutes at room temperature. A number of mAb concentrations were assessed to account for differences in binding strength between mAbs. Beads were subsequently washed and stained with 2.5 µg/mL goat anti-human IgG-AF647 secondary antibody (Jackson Immunoresearch). After a final wash, mAb-bound beads were read with the iQue Screener Plus (Intellicyt) high-throughput flow cytometer and data were analyzed with FlowJo (Tree Star).

### Growth inhibition activity (GIA) assay

The GIA was performed by the GIA Reference Center, NIH, USA as previously described (Miura et al., 2023). Briefly, all test antibodies were dialyzed against RPMI 1640. The indicated concentration of a test antibody was mixed with a trophozoite-rich 3D7 clone of *P. falciparum* in a 96-well plate (a total of 40 µL/well with 1% hematocrit at ∼0.3% parasitemia in the final well). The plate was incubated in an atmosphere of 5% O2, 5% CO2, and 90% N2 for ∼40 hours at 37°C. After the incubation, relative parasitemia in each well was quantified by parasite-specific lactate dehydrogenase (LDH) activity. The optical density at 650 nm (OD650) for each well was read by a VersaMax microplate reader and the % GIA value of each test sample was calculated as:

% GIA = 1-(OD650test – OD650uRBC) / (OD650iRBC – OD650uRBC) x 100

where OD650test, OD650uRBC, and OD650iRBC are average OD650 values for test antibody, uninfected RBC alone (no test antibody), and infected RBC alone (no test antibody) wells, respectively.

The antigen-reversal GIA was conducted as described elsewhere (Miura *et al*., 2007). Briefly, a test antibody was mixed with 0.5 mM of RH5 in a 96-well plate (RPMI 1640 was used as the buffer both for test antibody and RH5), then incubated for 45 minutes at room temperature followed by 15 minute incubation at 37°C. As done in the regular GIA described above, trophozoite-rich *P. falciparum* was added to the plate (a total of 40 µL/well with 1% hematocrit at

∼0.3% parasitemia in a final well), then the LDH assay was performed after a ∼40-hour incubation at 37°C.

### Crystallization and structure determination

To prepare complexes for crystallization, RH5ΔNL was treated with 1 μg/mL Endoproteinase Glu-C (Sigma) overnight at room temperature, then combined with scFv fragments of R5.008, MAD8-151, MAD8-502 or MAD10-255, or the Fab fragment of MAD10-466, in each case with RH5ΔNL in slight molar excess. Complexes were isolated by gel filtration using an S200 Increase 10/300 column running in SEC buffer (20 mM HEPES pH 7.5, 150 mM NaCl, 5% glycerol), then concentrated using 10K Amicon® Ultra centrifugal units. Complexes were crystallized by sitting drop vapour diffusion at 18°C mixing 100 nL each of protein and reservoir solution for all complexes except for R5.008 and MAD10-466, which are detailed below.

Crystals of RH5ΔNL:R5.008 were obtained at 7 mg/mL in Morpheus E11 (0.1 M Buffer System 3 pH 8.5, 0.12 M Ethylene Glycol Mix, 30% v/v Precipitant Mix 3) containing Silver Bullets G6 (0.02 M HEPES pH 0.16% w/v Glutaric acid, 0.16% Mellitic acid, 0.16% Oxalic acid anhydrous, 0.16% Pimelic acid, 0.16% Sebabic acid, 0.16% trans-Cinnamic acid), mixing 100 nL protein, 100 nL reservoir solution and 50 nL Silver Bullets additive. Crystals of RH5ΔNL:MAD10-466 were obtained at 7.5 mg/mL in Morpheus H9 (0.1 M Buffer System 3 pH 8.5, 0.1 M Amino Acids, 30% v/v Precipitant Mix 1) containing Hampton Additive G8 (40% v/v Polypropylene glycol P 400), mixing 100 nL protein, 50 nL reservoir solution, 50 nL additive, and 50 nL crystal seeds prepared using Seed Beads™ (Hampton Research) from crystals obtained in Morpheus H9 alone. Crystals of RH5ΔNL:MAD8-151 were obtained at 10 mg/mL in ProPlex E9 (0.2 M sodium chloride, 0.1 M sodium HEPES pH 7.5, 12% w/v PEG 8000), of RH5ΔNL:MAD8-502 at 11 mg/mL in JCSG+ D12 (0.04 M potassium phosphate monobasic, 16% w/v PEG 8000, 20% w/v glycerol), and of RH5ΔNL:MAD10-255 at 9 mg/mL in PGA H12 (0.1 M ammonium sulphate, 0.3 M sodium formate, 0.1 M Tris pH 7.8, 2% w/v γ-PGA (Na+ form, LM), 3% w/v PEG 20000). For data collection, crystals were cryoprotected where required in reservoir solution supplemented with 25% glycerol, then cryo-cooled in liquid nitrogen. Data were collected at Diamond Light Source on beamline I24 (wavelength 0.9999 Å) for RH5ΔNL:R5.008 and beamline I04 (wavelength 0.95374 Å) for all others.

Diffraction data for RH5ΔNL:R5.008 was auto-processed with xia2-multiplex combining two datasets acquired from the same crystal at different locations. Frames 1501-1800 of dataset 1 and frames 11501-11800 of dataset 2 were excluded due to radiation damage. The combined dataset was reduced, scaled, and merged using AIMLESS in CCP4i2 (Agirre et al., 2023) to 3.20 Å. Data for RH5ΔNL:MAD8-151 was auto-processed using autoPROC to 1.95 Å, and data for RH5ΔNL:MAD8-502, RH5ΔNL:MAD10-255 and RH5ΔNL:MAD10-466 were auto-processed in xia2-dials to 2.07 Å, 3.15 Å and 3.20 Å respectively. Molecular replacement for each dataset except RH5ΔNL:MAD10-466 was performed using PHASER in the CCP4 suite, using RH5 from PDB 6RCU and an scFv scaffold as search models. Molecular replacement of the MAD10-466 dataset was performed using the structure of RH5-bound MAD8-502 determined in this study and the constant domain of a Fab fragment (PDB ID 7PHU), also in PHASER. Models were built and refined in cycles using COOT (0.9.3) (Emsley et al., 2010), BUSTER (2.10.4) (Bricogne et al., 2017) and PHENIX (1.20.1-4487) (Liebschner et al., 2019). The unit cell for RH5ΔNL:R5.008 contains one copy of the RH5:scFv complex, while RH5ΔNL:MAD8-151 contains two copies, and RH5ΔNL:MAD8-502 contains four copies. RH5ΔNL:MAD10-255 contains five complete copies of the RH5:scFv complex and a further copy of RH5 alone with low occupancy for its partner scFv (which has therefore been omitted). The unit cell for RH5ΔNL:MAD10-466 contains two copies of the RH5:Fab fragment complex. For structural analysis, chains A and B for each RH5:scFv complex were used, or chains A, B and C for the RH5ΔNL:MAD10-466 Fab complex, with other chains in some cases showing weaker density. Binding interfaces were analyzed using PDBe PISA (Krissinel and Henrick, 2007), and structures were visualized and rendered in ChimeraX (Pettersen et al., 2021).

### Statistical analysis

Correlation r and P values for associations between antibody levels and age in the Malian cohort were generated by Spearman correlation. Comparisons between paired acute and convalescent samples were made by Wilcoxon sign-rank test. Comparisons between infection- and vaccination-derived mAbs were done by the Mann-Whitney U test. Spearman correlations for potency of mAbs in bins I and II with sequence and binding parameters was determined in R (*corrplot* package). r and P values for correlations between RH5 binding (AUC) and growth inhibition percentage were generated by Spearman correlation.

## REFERENCES

Agirre, J., Atanasova, M., Bagdonas, H., Ballard, C.B., Basle, A., Beilsten-Edmands, J., Borges, R.J., Brown, D.G., Burgos-Marmol, J.J., Berrisford, J.M., et al. (2023). The CCP4 suite: integrative software for macromolecular crystallography. Acta Crystallogr D Struct Biol 79, 449–461. 10.1107/S2059798323003595.

Alanine, D.G.W., Quinkert, D., Kumarasingha, R., Mehmood, S., Donnellan, F.R., Minkah, N.K., Dadonaite, B., Diouf, A., Galaway, F., Silk, S.E., et al. (2019). Human antibodies that slow erythrocyte invasion potentiate malaria-neutralizing antibodies. Cell 178, 216–228.e221.

Baum, J., Chen, L., Healer, J., Lopaticki, S., Boyle, M., Triglia, T., Ehlgen, F., Ralph, S.A., Beeson, J.G., and Cowman, A.F. (2009). Reticulocyte-binding protein homologue 5 - an essential adhesin involved in invasion of human erythrocytes by *Plasmodium falciparum*. Int J Parasitol 39, 371–380. 10.1016/j.ijpara.2008.10.006.

Bricogne, G., Blanc, E., Brandl, M., Flensburg, C., Keller, P., Paciorek, W., Roversi, P., Sharff, A., Smart, O.S., Vonrhein, C., and Womack, T.O. (2017). BUSTER version 2.10.4. Cambridge, United Kingdom: Global Phasing Ltd.

Bustamante, L.Y., Bartholdson, S.J., Crosnier, C., Campos, M.G., Wanaguru, M., Nguon, C., Kwiatkowski, D.P., Wright, G.J., and Rayner, J.C. (2013). A full-length recombinant *Plasmodium falciparum* PfRH5 protein induces inhibitory antibodies that are effective across common PfRH5 genetic variants. Vaccine 31, 373–379. 10.1016/j.vaccine.2012.10.106.

Chiu, C.Y., Healer, J., Thompson, J.K., Chen, L., Kaul, A., Savergave, L., Raghuwanshi, A., Li Wai Suen, C.S., Siba, P.M., Schofield, L., et al. (2014). Association of antibodies to *Plasmodium falciparum* reticulocyte binding protein homolog 5 with protection from clinical malaria. Front Microbiol 5, 314. 10.3389/fmicb.2014.00314.

Cho, H., Gonzales-Wartz, K.K., Huang, D., Yuan, M., Peterson, M., Liang, J., Beutler, N., Torres, J.L., Cong, Y., Postnikova, E., et al. (2021). Bispecific antibodies targeting distinct regions of the spike protein potently neutralize SARS-CoV-2 variants of concern. Sci Transl Med 13, eabj5413. 10.1126/scitranslmed.abj5413.

Crosnier, C., Bustamante, L.Y., Bartholdson, S.J., Bei, A.K., Theron, M., Uchikawa, M., Mboup, S., Ndir, O., Kwiatkowski, D.P., Duraisingh, M.T., et al. (2011). Basigin is a receptor essential for erythrocyte invasion by *Plasmodium falciparum*. Nature 480, 534–537. 10.1038/nature10606.

Crosnier, C., Wanaguru, M., McDade, B., Osier, F.H., Marsh, K., Rayner, J.C., and Wright, G.J. (2013). A library of functional recombinant cell-surface and secreted *P. falciparum* merozoite proteins. Mol Cell Proteomics 12, 3976–3986. 10.1074/mcp.O113.028357.

Dacon, C., Peng, L., Lin, T.H., Tucker, C., Lee, C.D., Cong, Y., Wang, L., Purser, L., Cooper, A.J.R., Williams, J.K., et al. (2023). Rare, convergent antibodies targeting the stem helix broadly neutralize diverse betacoronaviruses. Cell Host Microbe 31, 97–111 e112. 10.1016/j.chom.2022.10.010.

Datoo, M.S., Natama, H.M., Some, A., Bellamy, D., Traore, O., Rouamba, T., Tahita, M.C., Ido, N.F.A., Yameogo, P., Valia, D., et al. (2022). Efficacy and immunogenicity of R21/Matrix-M vaccine against clinical malaria after 2 years’ follow-up in children in Burkina Faso: a phase 1/2b randomised controlled trial. Lancet Infect Dis 22, 1728-1736. 10.1016/S1473-3099(22)00442-X.

Douglas, A.D., Baldeviano, G.C., Jin, J., Miura, K., Diouf, A., Zenonos, Z.A., Ventocilla, J.A., Silk, S.E., Marshall, J.M., Alanine, D.G.W., et al. (2019). A defined mechanistic correlate of protection against Plasmodium falciparum malaria in non-human primates. Nat Commun 10, 1953. 10.1038/s41467-019-09894-4.

Douglas, A.D., Baldeviano, G.C., Lucas, C.M., Lugo-Roman, L.A., Crosnier, C., Bartholdson, S.J., Diouf, A., Miura, K., Lambert, L.E., Ventocilla, J.A., et al. (2015). A PfRH5-based vaccine is efficacious against heterologous strain blood-stage *Plasmodium falciparum* infection in aotus monkeys. Cell Host Microbe 17, 130–139. 10.1016/j.chom.2014.11.017.

Douglas, A.D., Williams, A.R., Illingworth, J.J., Kamuyu, G., Biswas, S., Goodman, A.L., Wyllie, D.H., Crosnier, C., Miura, K., Wright, G.J., et al. (2011). The blood-stage malaria antigen PfRH5 is susceptible to vaccine-inducible cross-strain neutralizing antibody. Nat Commun 2, 601. 10.1038/ncomms1615.

Douglas, A.D., Williams, A.R., Knuepfer, E., Illingworth, J.J., Furze, J.M., Crosnier, C., Choudhary, P., Bustamante, L.Y., Zakutansky, S.E., Awuah, D.K., et al. (2014). Neutralization of *Plasmodium falciparum* merozoites by antibodies against PfRH5. J Immunol 192, 245–258. 10.4049/jimmunol.1302045.

Elton, C.M., Rodriguez, M., Ben Mamoun, C., Lobo, C.A., and Wright, G.J. (2019). A library of recombinant *Babesia microti* cell surface and secreted proteins for diagnostics discovery and reverse vaccinology. Int J Parasitol 49, 115–125. 10.1016/j.ijpara.2018.10.003.

Emsley, P., Lohkamp, B., Scott, W.G., and Cowtan, K. (2010). Features and development of Coot. Acta Crystallogr D Biol Crystallogr 66, 486–501. 10.1107/S0907444910007493.

Galaway, F., Yu, R., Constantinou, A., Prugnolle, F., and Wright, G.J. (2019). Resurrection of the ancestral RH5 invasion ligand provides a molecular explanation for the origin of *P. falciparum* malaria in humans. PLoS Biol 17, e3000490. 10.1371/journal.pbio.3000490.

Healer, J., Wong, W., Thompson, J.K., He, W., Birkinshaw, R.W., Miura, K., Long, C.A., Soroka, V., Sogaard, T.M.M., Jorgensen, T., et al. (2019). Neutralising antibodies block the function of Rh5/Ripr/CyRPA complex during invasion of *Plasmodium falciparum* into human erythrocytes. Cell Microbiol 21, e13030. 10.1111/cmi.13030.

Huang, J., Doria-Rose, N.A., Longo, N.S., Laub, L., Lin, C.L., Turk, E., Kang, B.H., Migueles, S.A., Bailer, R.T., Mascola, J.R., and Connors, M. (2013). Isolation of human monoclonal antibodies from peripheral blood B cells. Nat Protoc 8, 1907–1915. 10.1038/nprot.2013.117.

Jamwal, A., Constantin, C.E., Henrich, S., Bildl, W., Fakler, B., Draper, S.J., Schulte, U., and Higgins, M.K. (2022). Erythrocyte invasion-neutralising antibodies prevent *Plasmodium falciparum* RH5 from binding to basigin-containing membrane protein complexes. bioRxiv, 2022.2009.2023.509221. 10.1101/2022.09.23.509221.

Kayentao, K., Ongoiba, A., Preston, A.C., Healy, S.A., Doumbo, S., Doumtabe, D., Traore, A., Traore, H., Djiguiba, A., Li, S., et al. (2022). Safety and efficacy of a monoclonal antibody against malaria in Mali. N Engl J Med 387, 1833–1842. 10.1056/NEJMoa2206966.

Knuepfer, E., Wright, K.E., Kumar Prajapati, S., Rawlinson, T.A., Mohring, F., Koch, M., Lyth, O.R., Howell, S.A., Villasis, E., Snijders, A.P., et al. (2019). Divergent roles for the RH5 complex components, CyRPA and RIPR in human-infective malaria parasites. PLoS Pathog 15, e1007809. 10.1371/journal.ppat.1007809.

Krissinel, E., and Henrick, K. (2007). Inference of macromolecular assemblies from crystalline state. J Mol Biol 372, 774–797. 10.1016/j.jmb.2007.05.022.

Liebschner, D., Afonine, P.V., Baker, M.L., Bunkoczi, G., Chen, V.B., Croll, T.I., Hintze, B., Hung, L.W., Jain, S., McCoy, A.J., et al. (2019). Macromolecular structure determination using X-rays, neutrons and electrons: recent developments in Phenix. Acta Crystallogr D Struct Biol 75, 861–877. 10.1107/S2059798319011471.

Minassian, A.M., Silk, S.E., Barrett, J.R., Nielsen, C.M., Miura, K., Diouf, A., Loos, C., Fallon, J.K., Michell, A.R., White, M.T., et al. (2021). Reduced blood-stage malaria growth and immune correlates in humans following RH5 vaccination. Med (N Y) 2, 701–719 e719. 10.1016/j.medj.2021.03.014.

Miura, K., Diouf, A., Fay, M.P., Barrett, J.R., Payne, R.O., Olotu, A.I., Minassian, A.M., Silk, S.E., Draper, S.J., and Long, C.A. (2023). Assessment of precision in growth inhibition assay (GIA) using human anti-PfRH5 antibodies. Malar J 22, 159. 10.1186/s12936-023-04591-6.

Miura, K., Zhou, H., Muratova, O.V., Orcutt, A.C., Giersing, B., Miller, L.H., and Long, C.A. (2007). In immunization with *Plasmodium falciparum* apical membrane antigen 1, the specificity of antibodies depends on the species immunized. Infect Immun 75, 5827–5836. 10.1128/IAI.00593-07.

Moir, S., Lapointe, R., Malaspina, A., Ostrowski, M., Cole, C.E., Chun, T.W., Adelsberger, J., Baseler, M., Hwu, P., and Fauci, A.S. (1999). CD40-mediated induction of CD4 and CXCR4 on B lymphocytes correlates with restricted susceptibility to human immunodeficiency virus type 1 infection: potential role of B lymphocytes as a viral reservoir. J Virol 73, 7972–7980. 10.1128/JVI.73.10.7972-7980.1999.

Mwakingwe-Omari, A., Healy, S.A., Lane, J., Cook, D.M., Kalhori, S., Wyatt, C., Kolluri, A., Marte-Salcedo, O., Imeru, A., Nason, M., et al. (2021). Two chemoattenuated PfSPZ malaria vaccines induce sterile hepatic immunity. Nature 595, 289–294. 10.1038/s41586-021-03684-z.

Nouri, N., and Kleinstein, S.H. (2020). Somatic hypermutation analysis for improved identification of B cell clonal families from next-generation sequencing data. PLoS Comput Biol 16, e1007977. 10.1371/journal.pcbi.1007977.

Obeng-Adjei, N., Larremore, D.B., Turner, L., Ongoiba, A., Li, S., Doumbo, S., Yazew, T.B., Kayentao, K., Miller, L.H., Traore, B., et al. (2020). Longitudinal analysis of naturally acquired PfEMP1 CIDR domain variant antibodies identifies associations with malaria protection. JCI Insight 5. 10.1172/jci.insight.137262.

Partey, F.D., Castberg, F.C., Sarbah, E.W., Silk, S.E., Awandare, G.A., Draper, S.J., Opoku, N., Kweku, M., Ofori, M.F., Hviid, L., and Barfod, L. (2018). Kinetics of antibody responses to PfRH5-complex antigens in Ghanaian children with *Plasmodium falciparum* malaria. PLoS One 13, e0198371. 10.1371/journal.pone.0198371.

Pettersen, E.F., Goddard, T.D., Huang, C.C., Meng, E.C., Couch, G.S., Croll, T.I., Morris, J.H., and Ferrin, T.E. (2021). UCSF ChimeraX: Structure visualization for researchers, educators, and developers. Protein Sci 30, 70–82. 10.1002/pro.3943.

Ragotte, R.J., Higgins, M.K., and Draper, S.J. (2020). The RH5-CyRPA-Ripr complex as a malaria vaccine target. Trends in Parasitology 36, 545–559. 10.1016/j.pt.2020.04.003.

Ragotte, R.J., Pulido, D., Lias, A.M., Quinkert, D., Alanine, D.G.W., Jamwal, A., Davies, H., Nacer, A., Lowe, E.D., Grime, G.W., et al. (2022). Heterotypic interactions drive antibody synergy against a malaria vaccine candidate. Nat Commun 13, 933. 10.1038/s41467-022-28601-4.

Richards, J.S., Arumugam, T.U., Reiling, L., Healer, J., Hodder, A.N., Fowkes, F.J., Cross, N., Langer, C., Takeo, S., Uboldi, A.D., et al. (2013). Identification and prioritization of merozoite antigens as targets of protective human immunity to *Plasmodium falciparum* malaria for vaccine and biomarker development. J Immunol 191, 795–809. 10.4049/jimmunol.1300778.

Scally, S.W., Triglia, T., Evelyn, C., Seager, B.A., Pasternak, M., Lim, P.S., Healer, J., Geoghegan, N.D., Adair, A., Tham, W.H., et al. (2022). PCRCR complex is essential for invasion of human erythrocytes by *Plasmodium falciparum*. Nat Microbiol 7, 2039–2053. 10.1038/s41564-022-01261-2.

Silk, S.E., Kalinga, W.F., Mtaka, I.M., Lilolime, N.S., Mpina, M., Milando, F., Ahmed, S., Diouf, A., Mkwepu, F., Simon, B., et al. (2023). Superior antibody immunogenicity of a viral-vectored RH5 blood-stage malaria vaccine in Tanzanian infants as compared to adults. Med. 10.1016/j.medj.2023.07.003.

Takala, S.L., Coulibaly, D., Thera, M.A., Batchelor, A.H., Cummings, M.P., Escalante, A.A., Ouattara, A., Traore, K., Niangaly, A., Djimde, A.A., et al. (2009). Extreme polymorphism in a vaccine antigen and risk of clinical malaria: implications for vaccine development. Sci Transl Med 1, 2ra5. 10.1126/scitranslmed.3000257.

Tan, J., Cho, H., Pholcharee, T., Pereira, L.S., Doumbo, S., Doumtabe, D., Flynn, B.J., Schon, A., Kanatani, S., Aylor, S.O., et al. (2021). Functional human IgA targets a conserved site on malaria sporozoites. Sci Transl Med 13. 10.1126/scitranslmed.abg2344.

Tran, T.M., Guha, R., Portugal, S., Skinner, J., Ongoiba, A., Bhardwaj, J., Jones, M., Moebius, J., Venepally, P., Doumbo, S., et al. (2019). A molecular signature in blood reveals a role for p53 in regulating malaria-induced inflammation. Immunity 51, 750–765 e710. 10.1016/j.immuni.2019.08.009.

Tran, T.M., Ongoiba, A., Coursen, J., Crosnier, C., Diouf, A., Huang, C.Y., Li, S., Doumbo, S., Doumtabe, D., Kone, Y., et al. (2014). Naturally acquired antibodies specific for *Plasmodium falciparum* reticulocyte-binding protein homologue 5 inhibit parasite growth and predict protection from malaria. J Infect Dis 209, 789–798. 10.1093/infdis/jit553.

Wang, L.T., Pereira, L.S., Flores-Garcia, Y., O’Connor, J., Flynn, B.J., Schon, A., Hurlburt, N.K., Dillon, M., Yang, A.S.P., Fabra-Garcia, A., et al. (2020). A potent anti-malarial human monoclonal antibody targets circumsporozoite protein minor repeats and neutralizes sporozoites in the liver. Immunity 53, 733–744.e738. 10.1016/j.immuni.2020.08.014.

Willcox, A.C., Huber, A.S., Diouf, A., Barrett, J.R., Silk, S.E., Pulido, D., King, L.D.W., Alanine, D.G.W., Minassian, A.M., Diakite, M., et al. (2021). Antibodies from malaria-exposed Malians generally interact additively or synergistically with human vaccine-induced RH5 antibodies. Cell Rep Med 2, 100326. 10.1016/j.xcrm.2021.100326.

Wong, W., Huang, R., Menant, S., Hong, C., Sandow, J.J., Birkinshaw, R.W., Healer, J., Hodder, A.N., Kanjee, U., Tonkin, C.J., et al. (2019). Structure of *Plasmodium falciparum* Rh5-CyRPA-Ripr invasion complex. Nature 565, 118–121. 10.1038/s41586-018-0779-6.

Wright, G.J., and Rayner, J.C. (2014). *Plasmodium falciparum* erythrocyte invasion: combining function with immune evasion. PLoS Pathog 10, e1003943. 10.1371/journal.ppat.1003943.

Wright, K.E., Hjerrild, K.A., Bartlett, J., Douglas, A.D., Jin, J., Brown, R.E., Illingworth, J.J., Ashfield, R., Clemmensen, S.B., de Jongh, W.A., et al. (2014). Structure of malaria invasion protein RH5 with erythrocyte basigin and blocking antibodies. Nature 515, 427–430. 10.1038/nature13715.

Wu, R.L., Idris, A.H., Berkowitz, N.M., Happe, M., Gaudinski, M.R., Buettner, C., Strom, L., Awan, S.F., Holman, L.A., Mendoza, F., et al. (2022). Low-dose subcutaneous or intravenous monoclonal antibody to prevent malaria. N Engl J Med 387, 397–407. 10.1056/NEJMoa2203067.

Wu, X., Yang, Z.Y., Li, Y., Hogerkorp, C.M., Schief, W.R., Seaman, M.S., Zhou, T., Schmidt, S.D., Wu, L., Xu, L., et al. (2010). Rational design of envelope identifies broadly neutralizing human monoclonal antibodies to HIV-1. Science 329, 856-861. 10.1126/science.1187659.

Zenonos, Z.A., Rayner, J.C., and Wright, G.J. (2014). Towards a comprehensive *Plasmodium falciparum* merozoite cell surface and secreted recombinant protein library. Malar J 13, 93. 10.1186/1475-2875-13-93.

