## Supplemental Files for "Natural malaria infection elicits rare but potent neutralizing antibodies to the blood-stage antigen RH5"

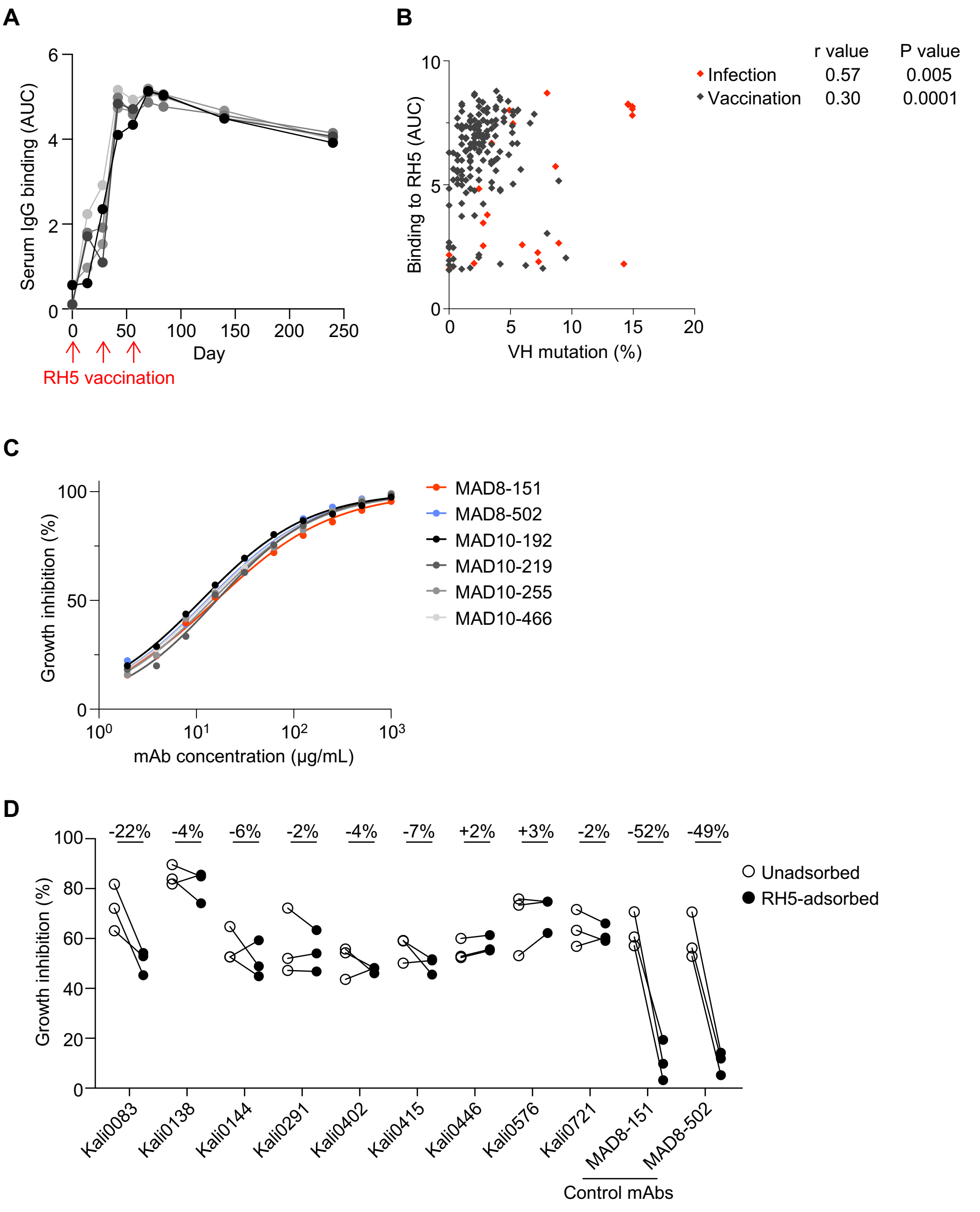


**Figure S1. Infection- and vaccination-induced antibody responses to RH5.**

A. Serum IgG reactivity to RH5 after three doses of the RH5/AS01 vaccine in malaria-naïve individuals.

B. Association between RH5 binding and VH mutations of RH5-specific mAbs from natural infection (red) and vaccination (black). P and r values were calculated based on Spearman correlation.

C. Growth inhibition titration curves of the six most potent RH5-specific mAbs. Data are shown from a representative experiment out of n = 2-3 experiments. MAD8-151 and MAD8-502 were isolated from infected donors while MAD10-192, MAD10-219, MAD10-255 and MAD10-466 were isolated from vaccinated donors.

D. Growth inhibition mediated by polyclonal IgG from naturally infected donors in an antigen-reversal assay, where samples are tested for activity with and without adsorption of RH5-specific antibodies with soluble antigen. Each pair of points represents an independent experiment. MAD8-151 and MAD8-502 are control RH5-specific mAbs. Kali0083 is the source donor of MAD8-502 and Kali0446 is the source donor of MAD8-151. The percentages at the top of the figure refer to the mean difference between the RH5-adsorbed and unadsorbed inhibition values.

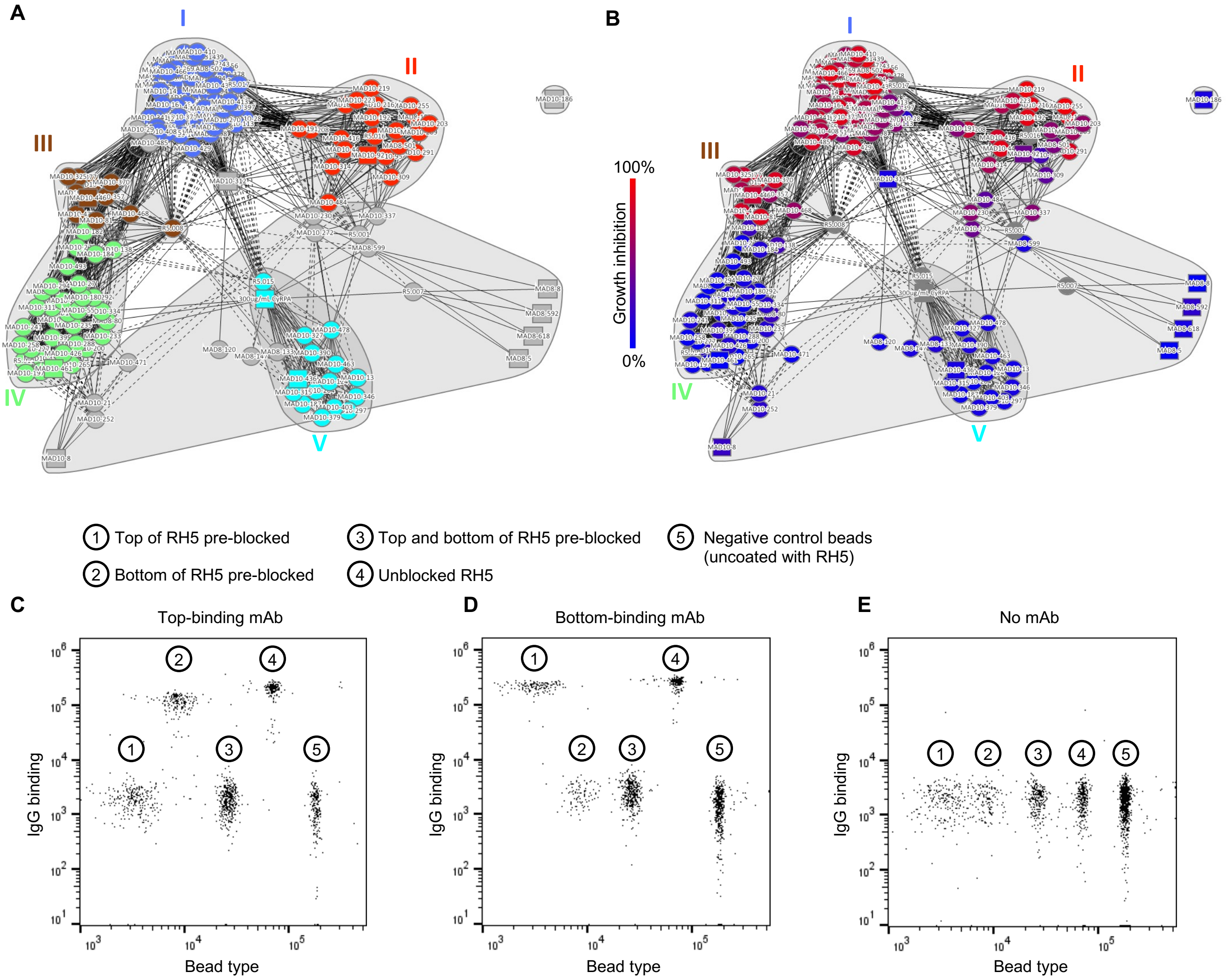


**Figure S2: Discrete RH5 epitope communities correlate with antibody GIA score.**

A. Epitope bins of RH5-specific mAbs clustered based on competition profiles. Solid lines indicate two-way competition, dashed lines indicate one-way competition. Circular nodes represent mAbs analyzed as both ligand and analyte; square nodes represent mAbs analyzed as ligand or analyte only.

B. GIA-coded epitope bins of RH5-specific mAbs. Nodes are color-coded by GIA percentage at 1 mg/mL. Negative scores were set to zero. Reference mAbs and the antigen CyRPA, which were not analyzed by GIA, are colored grey. Solid lines indicate two-way competition, dashed lines indicate one-way competition. Circular nodes represent mAbs analyzed as both ligand and analyte; square nodes represent mAbs analyzed as ligand or analyte only.

C-E. Representative FACS plots from a multiplex bead-based assay to determine epitope localization of RH5-specific mAbs. Bead populations 1-3 display RH5 that was pre-blocked at top epitopes, bottom epitopes, or both. Bead population 4 displays unblocked RH5. Bead population 5 was not coated with RH5. Fluorescently labeled anti-IgG secondary antibody was used to determine the level of binding of each mAb. Results from a representative top-binding mAb (C), bottom-binding mAb (D), and no mAb negative control (E) are shown.

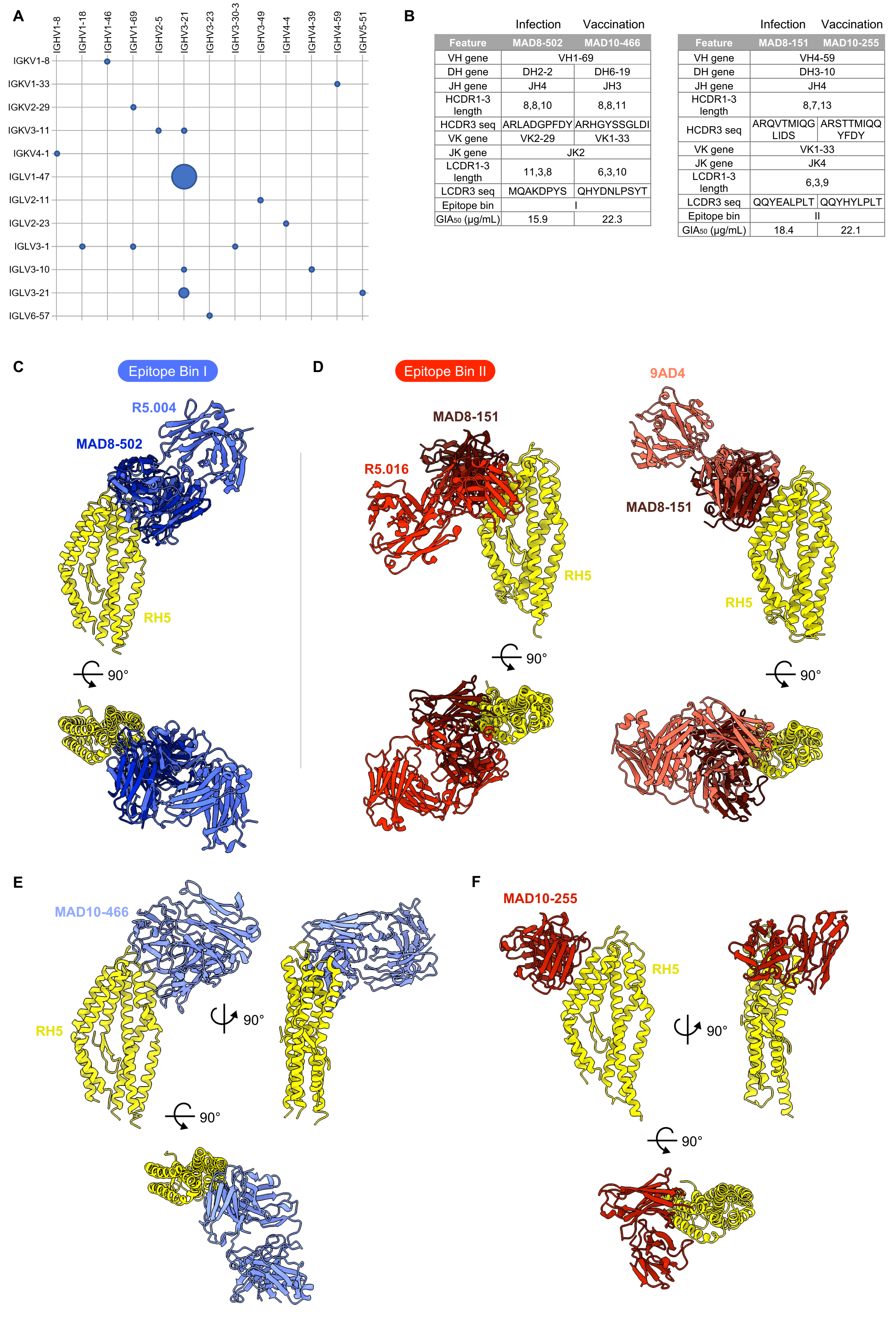


| Feature | MAD8-502 (Infection) | MAD10-466 (Vaccination) |
|---|---|---|
| VH gene | VH1-69 | |
| DH gene | DH2-2 | DH6-19 |
| JH gene | JH4 | JH3 |
| HCDR1-3 length | 8,8,10 | 8,8,11 |
| HCDR3 seq | ARLADGPFDY | ARHGYSSGLDI |
| VK gene | VK2-29 | VK1-33 |
| JK gene | JK2 | |
| LCDR1-3 length | 11,3,8 | 6,3,10 |
| LCDR3 seq | MQAKDPYS | QHYDNLPSYT |
| Epitope bin | I | |
| $GIA_{50}$ (μg/mL) | 15.9 | 22.3 |

| Feature | MAD8-151 (Infection) | MAD10-255 (Vaccination) |
|---|---|---|
| VH gene | VH4-59 | |
| DH gene | DH3-10 | |
| JH gene | JH4 | |
| HCDR1-3 length | 8,7,13 | |
| HCDR3 seq | ARQVTMIQG LIDS | ARSTTMIQQ YFDY |
| VK gene | VK1-33 | |
| JK gene | JK4 | |
| LCDR1-3 length | 6,3,9 | |
| LCDR3 seq | QQYEALPLT | QQYHYLPLT |
| Epitope bin | II | |
| $GIA_{50}$ (μg/mL) | 18.4 | 22.1 |

**Figure S3: Genetic and structural features of RH5-specific antibodies from infection and vaccination.**

A. Heavy (x-axis) and light (y-axis) chain V gene usage of RH5-specific mAbs from natural infection. The diameters of the circles are proportional to the number of mAbs using the corresponding heavy and light chain pair. The five mAbs using IGHV3-21/IGLV1-47 are clonally related, while the two mAbs using IGHV3-21/IGLV3-10 are not related and were isolated from different donors.

B. Genetic, binding and functional features of MAD8-502 in comparison to MAD10-466, and MAD8-151 in comparison to MAD10-255.

C. Structural alignments of MAD8-502 scFv (dark blue) and R5.004 Fab (blue, PDB ID 6RCU) bound to RH5 (yellow), showing that MAD8-502 belongs to epitope bin I. Complexes are shown as side and top views.

D. Structural alignments of MAD8-151 scFv (dark red) and R5.016 Fab (red, PDB ID 6RCV) or 9AD4 Fab (pink, PDB ID 4U0R) bound to RH5 (yellow), showing that MAD8-151 belongs to epitope bin II and shares an angle of approach to RH5 more similar to 9AD4 than R5.016.

E. Top and side views of the crystal structure of MAD10-466 Fab fragment (light blue) bound to RH5 (yellow).

F. Top and side views of the crystal structure of MAD10-255 scFv (red) bound to RH5 (yellow).

### Table S1. Crystallographic data collection and refinement statistics

| | RH5ΔNL:R5.008 | RH5ΔNL:MAD8-151 | RH5ΔNL:MAD8-502 | RH5ΔNL:MAD10-255 | RH5ΔNL:MAD10-466 |
|---|---|---|---|---|---|
| **Data Collection** | | | | | |
| Space group | P 31 2 1 | P 1 21 1 | P 1 21 1 | P1 21 1 | P 1 21 1 |
| Cell dimensions | | | | | |
| a, b, c (Å) | 114.99, 114.99, 147.49 | 78.30, 121.38, 79.00 | 85.97, 158.01, 113.02 | 115.31, 134.52, 198.67 | 82.46, 140.41, 83.62 |
| $\alpha, \beta, \gamma$ (°) | 90, 90, 120 | 90, 90.18, 90 | 90, 94.59, 90 | 90, 93.56, 90 | 90, 112.74, 90 |
| Wavelength | 0.9999 Å | 0.95374 | 0.95374 | 0.95374 | 0.95374 |
| Resolution (Å) | 57.49 – 3.20 (3.42-3.20) | 79.00-1.95 (1.98-1.95) | 79.01-2.07 (2.11-2.07) | 87.45-3.15 (3.20-3.15) | 67.59-3.20 (3.26-3.20) |
| Total Observations | 306591 (55698) | 730651 (30772) | 1245190 (41161) | 739821 (34690) | 205662 (10293) |
| Total Unique | 19138 (3408) | 106577 (5404) | 181986 (8654) | 104997 (5215) | 28799 (1404) |
| $R_{merge}$ (%) | 49.6 (301.0) | 25.2 (397.6) | 12.6 (328.2) | 19.0 (231.3) | 55.1 (291.3) |
| $R_{meas}$ (%) | 51.4 (311.4) | 27.3 (437.3) | 13.6 (369.7) | 20.5 (251.0) | 59.4 (313.5) |
| $R_{pim}$ (%) | 13.2 (78.3) | 10.4 (180.2) | 5.2 (166.4) | 7.7 (97.0) | 22.1 (115.4) |
| $CC_{1/2}$ | 0.986 (0.732) | 0.993 (0.337) | 0.997 (0.370) | 0.996 (0.264) | 0.952 (0.345) |
| $I/\sigma(I)$ | 5.2 (1.0) | 6.0 (0.7) | 7.5 (0.2) | 7.3 (0.4) | 2.3 (0.3) |
| Completeness (%) | 100.00 (99.9) | 98.5 (100.0) | 99.7 (95.0) | 100.0 (99.1) | 99.3 (97.0) |
| Multiplicity | 16.00 (16.3) | 6.9 (5.7) | 6.8 (4.8) | 7.0 (6.7) | 7.1 (7.3) |
| Wilson B factor | 70.8 | 27.9 | 49.4 | 102.5 | 74.0 |
| **Refinement** | | | | | |
| Reflections | 19053 | 105428 | 175452 | 104799 | 28644 |
| Rwork / Rfree (%) | 24.02/29.35 | 23.01/25.15 | 26.87/27.91 | 24.19/26.36 | 27.19/31.67 |
| Average B factor | | | | | |
| *Protein (all)* | 90.4 | 32.6 | 72.2 | 136.0 | 100.89 |
| *Protein (Chain A)* | 82.4 | 34.4 | 75.3 | 115.7 | 94.53 |
| *Protein (Chain B)* | 96.2 | 29.5 | 61.0 | 106.0 | 90.31 |
| *Protein (Chain C)* | - | - | - | - | 20.07 |
| *Water* | 74.0 | 51.6 | 63.6 | n/a | n/a |
| *Ligands* | 70.7 | 37.3 | n/a | n/a | n/a |
| Number of residues | | | | | |
| *Protein* | 516 | 1034 | 2038 | 2848 | 1383 |
| *Water* | 6 | 533 | 418 | 0 | 0 |
| *Ligands* | 1 | 7 | 0 | 0 | 0 |
| RMSDs | | | | | |
| *Bond lengths (Å)* | 0.008 | 0.008 | 0.008 | 0.009 | 0.007 |
| *Bond angles (°)* | 0.97 | 0.95 | 0.98 | 1.03 | 0.92 |
| Ramachandran plot | | | | | |
| *Favored (%)* | 92.0 | 97.8 | 97.9 | 93.2 | 92.1 |
| *Allowed (%)* | 8.0 | 2.0 | 2.1 | 6.8 | 7.9 |
| *Outliers (%)* | 0.0 | 0.2 | 0.0 | 0.0 | 0.0 |

**Table S2. Table of contacts between R5.008, MAD8-502, MAD10-466, MAD8-151 and MAD10-255 with RH5.**

| R5.008 | | | | | | | |
|---|---|---|---|---|---|---|---|
| Chain | Residue | Group | Location | Chain | Residue | Group | Interaction |
| A | D59 | Side chain | CDR H2 | B | K457 | Side chain | Salt bridge |
| A | Y103 | Side chain | CDR H3 | B | D440 | Side chain | H bond |
| A | Y104 | Side chain | CDR H3 | B | N445 | Side chain | H bond |
| | | Side chain | CDR H3 | B | D461 | Side chain | H bond |
| | | Side chain | CDR H3 | B | R458 | Side chain | Cation-$\pi$ |
| A | S106 | Side chain | CDR H3 | B | R458 | Side chain | H bond |
| A | Q164 | Side chain | CDR L1 | B | N349 | Side chain | H bond |
| A | Y169 | Side chain | CDR L1 | B | R448 | Side chain | H bond |
| A | Y229 | Backbone | CDR L3 | B | R448 | Side chain | H bond |
| | | Side chain | CDR L3 | B | T449 | Side chain | H bond |
| | | Side chain | CDR L3 | B | N349 | Backbone | H bond |
| A | T231 | Side chain | CDR L3 | B | D453 | Side chain | H bond |

| MAD8-502 | | | | | | | |
|---|---|---|---|---|---|---|---|
| Chain | Residue | Group | Location | Chain | Residue | Group | Interaction |
| B | T28 | Side chain | CDR H1 | A | H365 | Side chain | H bond |
| B | S31 | Side chain | CDR H1 | A | D361 | Side chain | H bond |
| | | Backbone | CDR H1 | | | Side chain | H bond |
| | | Backbone | CDR H1 | A | W447 | Side chain | H bond |
| B | I52 | Side chain | CDR H2 | | | | Hydrophobic |
| B | I54 | Side chain | CDR H2 | | | | Hydrophobic |
| B | L55 | Side chain | CDR H2 | | | | Hydrophobic |
| B | R57 | Side chain | CDR H2 | A | W447 | Backbone | H bond |
| B | D101 | Side chain | CDR H3 | A | Y358 | Side chain | H bond |
| | | Side chain | CDR H3 | A | R357 | Side chain | Salt bridge |
| B | P103 | Side chain | CDR H3 | | | | Hydrophobic |
| B | F104 | Side chain | CDR H3 | | | | Hydrophobic |
| B | S164 | Backbone | CDR L1 | A | N347 | Side chain | H bond |
| B | D165 | Side chain | CDR L1 | A | N352 | Side chain | H bond |
| | | | | A | Y358 | Side chain | Hydrophobic |
| | | | | A | K443 | Side chain | Hydrophobic |
| | | | | A | W447 | Side chain | Hydrophobic |
| | | | | A | I446 | Side chain | Hydrophobic |
| | | | | A | I442 | Side chain | Hydrophobic |

| MAD10-466 | | | | | | | |
|---|---|---|---|---|---|---|---|
| Chain | Residue | Group | Location | Chain | Residue | Group | Interaction |
| B | T28 | Side chain | CDR H1 | A | H365 | Side chain | H bond |
| B | S31 | Side chain | CDR H1 | A | D361 | Side chain | H bond |
| | S31 | Backbone | CDR H1 | A | D361 | Side chain | H bond |
| | S31 | Backbone | CDR H1 | A | R357 | Side chain | H bond |
| B | S31 | Backbone | CDR H1 | A | W447 | Side chain | H bond |
| B | I52 | Side chain | CDR H2 | | | | Hydrophobic |
| B | I54 | Backbone | CDR H2 | A | K443 | Side chain | H bond |
| | I54 | Side chain | CDR H2 | | | | Hydrophobic |
| B | F55 | Side chain | CDR H2 | | | | Hydrophobic |
| B | H99 | Backbone | CDR H3 | A | R357 | Side chain | H bond |
| B | G100 | Backbone | CDR H3 | A | R357 | Side chain | H bond |
| B | Y101 | Side chain | CDR H3 | A | D361 | Side chain | H bond |
| | Y101 | Backbone | CDR H3 | A | N354 | Side chain | H bond |
| | Y101 | Backbone | CDR H3 | | | | Hydrophobic |
| B | S103 | Side chain | CDR H3 | A | N352 | Side chain | H bond |
| | S103 | Side chain | CDR H3 | A | N354 | Side chain | H bond |
| C | D50 | Side chain | CDR L2 | A | N352 | Side chain | H bond |
| C | N53 | Side chain | CDR L2 | A | S197 | Side chain | H bond |

| | | | | | | | |
|---|---|---|---|---|---|---|---|
| C | S67 | Side chain | FWR 3 | A | Y346 | Side chain | H bond |
| C | N93 | Side chain | CDR L3 | A | W447 | Backbone | H bond |
| | | | | A | I442 | Side chain | Hydrophobic |
| | | | | A | I446 | Side chain | Hydrophobic |
| | | | | A | W447 | Side chain | Hydrophobic |

| MAD8-151 | | | | | | | |
|---|---|---|---|---|---|---|---|
| **Chain** | **Residue** | **Group** | **Location** | **Chain** | **Residue †** | **Group** | **Interaction** |
| B | S31 | Side chain | CDR H1 | A | N334 (N285) | Side chain | H bond |
| | | Side chain | CDR H1 | A | N334 (N285) | Backbone | H bond |
| B | Y32 | Side chain | CDR H1 | A | T337 (T288) | Side chain | H bond |
| | | Side chain | CDR H1 | A | E341 (E292) | Side chain | H bond |
| B | Y33 | Side chain | CDR H1 | A | D331 (D282) | Side chain | H bond |
| B | Y52 | Side chain | CDR H2 | A | K327 (K278) | Side chain | H bond |
| B | H53 | Side chain | CDR H2 | A | D331 (D282) | Side chain | Salt bridge |
| B | R97 | Side chain | CDR H3 | A | N338 (N289) | Side chain | H bond |
| B | V99 | Backbone | CDR H3 | A | N334 (N285) | Side chain | H bond |
| B | M101 | Backbone | CDR H3 | A | D331 (D282) | Side chain | H bond |
| B | I102 | Backbone | CDR H3 | A | D331 (D282) | Side chain | H bond |
| | | Side chain | CDR H3 | | | | Hydrophobic |
| B | D184 | Side chain | CDR L2 | A | K212 (K212) | Side chain | Salt bridge |
| B | Y225 | Side chain | CDR L3 | A | K212 (K212) | Side chain | H bond |
| | | | | A | F209 (F209) | Side chain | Hydrophobic |
| | | | | A | I213 (I213) | Side chain | Hydrophobic |
| | | | | A | A216 (A216) | Side chain | Hydrophobic |
| | | | | A | I328 (I279) | Side chain | Hydrophobic |

| MAD10-255 | | | | | | | |
|---|---|---|---|---|---|---|---|
| **Chain** | **Residue** | **Group** | **Location** | **Chain** | **Residue** | **Group** | **Interaction** |
| B | T31 | Side chain | CDR H1 | A | N334 | Side chain | H bond |
| | | Backbone | CDR H1 | A | N334 | Side chain | H bond |
| | | Side chain | CDR H1 | A | K333 | Side chain | H bond |
| B | Y32 | Side chain | CDR H1 | A | E341 | Side chain | H bond |
| B | Y33 | Side chain | CDR H1 | A | D331 | Side chain | H bond |
| B | H53 | Side chain | CDR H2 | A | D331 | Side chain | Salt bridge |
| B | R97 | Side chain | CDR H3 | A | N338 | Side chain | H bond |
| B | T99 | Backbone | CDR H3 | A | N334 | Side chain | H bond |
| B | M101 | Backbone | CDR H3 | A | D331 | Side chain | H bond |
| B | I102 | Backbone | CDR H3 | A | D331 | Side chain | H bond |
| | | Backbone | CDR H3 | A | K212 | Side chain | H bond |
| | | Side chain | CDR H3 | | | | Hydrophobic |
| B | D107 | Side chain | CDR H3 | A | N338 | Side chain | H bond |
| B | D184 | Side chain | CDR L2 | A | K212 | Side chain | Salt Bridge |
| B | Y225 | Side chain | CDR L3 | A | K212 | Side chain | H bond |
| | | | | A | F209 | Side chain | Hydrophobic |
| | | | | A | I213 | Side chain | Hydrophobic |
| | | | | A | A216 | Side chain | Hydrophobic |
| | | | | A | I328 | Side chain | Hydrophobic |

† note that for the structure of PfRH5 bound to MAD8-151, the PDB is numbered according to the PfRH5ΔNL numbering, as the linker replacing the deleted loop is observed. The numbers given here are those for full-length PfRH5, to match the numbering in other structures, while the numbers in paratheses are those in the PDB, which match numbering for PfRH5ΔNL.
